# Parental age effects on offspring telomere length across vertebrates: a meta-analysis

**DOI:** 10.1101/2025.07.26.666864

**Authors:** Mariia Vlasova, Yuheng Sun, Heung Ying Janet Chik, Hannah L. Dugdale

**Affiliations:** Groningen Institute for Evolutionary Life Sciences, University of Groningen, Linnaeusborg, Groningen, the Netherlands; School of Natural Sciences, Faculty of Science and Engineering, Macquarie University, Sydney, New South Wales, Australia

**Keywords:** aging, intergenerational effects, maternal age at conception, meta-analysis, paternal age at conception, senescence, systematic review, telomere length

## Abstract

Telomeres shorten with advancing age in numerous species, and shorter telomeres are linked to increased mortality risk. While parental age at conception can influence offspring telomere length, the magnitude and direction of this effect differ across studies, species, and parental sexes. A significant knowledge gap therefore exists in understanding how parental age influences offspring telomere length across vertebrates. To address this knowledge gap, we conducted a systematic review and meta-analysis to examine the effects of paternal and maternal age at conception on offspring telomere length, incorporating 99 effect sizes from 30 human studies and 49 effect sizes from 12 non-human vertebrates. Within the human studies, there was a positive overall parental age effect on offspring telomere length, while no effect was found in the non-human vertebrate studies after adjusting for study, estimate, and phylogenetic effects. There was considerable heterogeneity in both human and non-human studies. In human studies, heterogeneity was mainly attributed to between-study variance, and in non-human studies to phylogeny. Parental age effect estimates were correlated with the laboratory methods used for measuring telomere length in all studies. In human studies, the interaction between parental sex and offspring sex, and the cell type used for telomere extraction affected the parental age effect estimates. In non-human vertebrates, the parental age effects were more positive when the parents’ identity was controlled for in the study. We recommend that future research be conducted on a broader range of taxa, test for within-parent effects, and follow standardized reporting practices to enhance data comparability.

## Introduction

Telomere length (TL) is a biomarker of body state that has been drawing increasing attention within ecology and evolution biology (Monaghan et al., 2022; Wilbourn et al., 2018). Telomeres are complexes composed of proteins and nucleotides in tandem repeats (typically TTAGGG in vertebrates) at the ends of eukaryotic chromosomes (Monaghan & Haussmann, 2006; von Zglinicki, 2002). They act as protective chromosome caps, ensuring the integrity of genetic information during cell division (Bernadotte et al., 2016; López-Otín et al., 2013). Telomeres undergo progressive shortening due to the end-replication problem with each cell division (Levy et al., 1992); or due to oxidative damage (Houben et al., 2008), and their lengths are considered as a biomarker of senescence and/or oxidative stress (Houben et al., 2008; Mather et al., 2011). Telomeres shorten with increasing age in many species, and shorter telomeres are associated with higher mortality risk (Remot et al., 2021; Wilbourn et al., 2018). Telomeres can also be lengthened with the presence of active telomerase in some species, life stages and cell types (Bateson & Nettle, 2017; Cesare & Reddel, 2010; Gomes et al., 2011). Telomere dynamics are influenced by factors ranging from inherited determinants, parental age, prenatal conditions, early life development and telomerase activity (reviewed in (Monaghan et al., 2018).

Testing for factors that determine TL helps us better understand the ecology of senescence (the decline in performance and fitness in later life). Parents can affect offspring phenotypes (including offspring lifespan and fitness-related traits), beyond the DNA that the offspring directly inherit from their parents, via both genetic and non-genetic pathways such as parental care, and these effects are collectively known as parental effects (Ivimey-Cook et al., 2022; Monaghan et al., 2020).

Parental effects, and in particular, parental age at conception (Starkweather et al., 2014) impacts offspring TL and its subsequent dynamics, but the magnitude and direction of this effect varies across studies, species, and parental sexes. In humans, there is evidence of a positive association between offspring TL and paternal age (Arbeev et al., 2011; Broer et al., 2013), but a negative or absent association for maternal age (Keefe et al., 2007). In non-human vertebrates, there is evidence for a negative correlation between paternal age and offspring TL (Criscuolo et al., 2017; McLennan et al., 2018), a negative (Carnes et al., 2011) or positive (Reichert et al., 2014) correlation between maternal age and offspring TL, or no significant parental age correlation at all (Belmaker et al., 2019; Froy et al., 2017).

These inconsistent parental age effects on TL can come from several sources. First, different maternal and paternal age effects can be due to the differences in female and male reproductive biology. In females, eggs ovulated in later life undergo more cell replications before they enter meiosis during egg formation in embryogenesis than eggs ovulated in earlier life, leading to shorter telomeres (Keefe, 2016). Also, oocytes remain in the ovary until ovulation, so eggs ovulated later in life are exposed to more oxidative stress, which contributes to shorter telomeres compared to eggs ovulated in earlier life (Keefe, 2016). Altogether, offspring produced in later life may therefore inherit shorter telomeres. In contrast, males produce germ cells continuously throughout their reproductive lives, and telomerase, which maintains and lengthens telomeres, is active in the relevant stem cell population across adult life at high levels in human testes (Eisenberg & Kuzawa, 2018; Wright et al., 1996). Moreover, the selective loss of spermatogonial stem cells with short telomeres can shift the distribution of sperm telomeres, resulting in longer gamete telomeres in older fathers (Eisenberg & Kuzawa, 2018). As a result, offspring TL can increase with paternal age.

Second, the parental age effects on offspring TL may also depend on the sex of the offspring. Advancing parental age can lead to shorter offspring TL because of a reduction in the quality of parental care with age, and the amount of parental care may affect sons and daughters differently (Marasco et al., 2019; Monaghan et al., 2020). This may be caused by sons and daughters responding differently to their early-life environments or through sex-specific epigenetic inheritance mechanisms (Bouwhuis et al., 2015; Schroeder et al., 2015). As a result, the decline in parental care quality with age may cause sex-specific telomere shortening (Marasco et al., 2019). Moreover, parental age effects may occur only with certain parent-offspring sex combinations. For example, in wild house sparrows (*Passer domesticus*) and common terns (*Sterna hirundo*), daughters born to older mothers had lower fitness than her earlier-born sisters, and the same for sons of older fathers, and it is hypothesised that telomere shortening mediated these intergenerational effects (Bouwhuis et al., 2015; Schroeder et al., 2015).

Third, the parental age effects on offspring TL can differ when they are measured cross-sectionally (between parents) than when they are measured longitudinally (within parents). A positive association or no association between parental age and offspring TL can be observed between-parents due to selective disappearance, i.e. only parents with long telomeres survive to old age and they produce offspring with long telomeres (Monaghan, 2024). However, if measured repeatedly within a parent, it is still possible to observe a negative association when there is a positive or no association cross-sectionally: as a parent ages the offspring it produces in later life have shorter telomeres than the offspring it produces in earlier life (Sparks et al., 2022). Longitudinal studies allow the separation of the between-parent and within-parent effects (van de Pol & Wright, 2009). Briefly, in these longitudinal studies, the mean age per parent at conception captures the between-parent age effect, and the difference from the mean age at conception of the parent captures the within-parent age effect. This method has been applied in several empirical studies of parental age effects on TL (e.g. (Sparks et al., 2021; van Lieshout et al., 2021).

Other factors can also contribute to the complexity of parental age effects. For example, (1) telomere biology and parental effects vary across species (Gomes et al., 2011; Ivimey-Cook et al., 2022), which can lead to variation in parental age effects on offspring TL between species. (2) Since the paternal age effect on TL is thought to be driven by continual production of sperm, the paternal age effect on offspring TL in species that produce sperm for a small part of the year may be less positive/more negative compared with species that produce sperm year-round (Eisenberg & Kuzawa, 2018). (3) Methodological and qualitative differences in TL extraction and assessment methods can also cause differences, mostly among studies. For example, TL measured by real-time quantitative polymerase chain reaction (qPCR) has larger variation than those measured by the terminal restriction fragments (TRF) method, and they are expressed in different units which leads to incomparability (Aviv, 2008; Verhulst, 2019). Furthermore, TL measured with TRF methods using Southern blots is longer than those measured with in-gel hybridization techniques because the latter exclude interstitial telomeres which occur in some species (Bolzán, 2017; Nussey et al., 2014). Whole genome sequencing (WGS) based techniques provide an alternative to TL measuring techniques when WGS data are available (Lee et al., 2017). WGS-based measurements correlate well with qPCR measurements, but not with TRF analysis, and they can filter out interstitial telomeric sequences using different approaches (Lee et al., 2017) (4) Telomere shortening rates vary across tissues, thus the type of tissue in which TL was sampled may affect the results (McLester-Davis et al., 2023; Wilbur et al., 2019). (5) Parental environment, especially adverse maternal environments are associated with shorter offspring TL, so the parental environment effect may bias the parental age effect measured (Lulkiewicz et al., 2020). (6) Whether the parental identity was accounted for may influence effect size estimates, as not controlling for repeated measures from the same parents might lead to non-independent data points, which can inflate significance (Hurlbert, 1984). (7) Offspring age at TL measurement can also play a role, because the offspring can have the same initial TLs, but TL declines with adult age (Remot et al., 2021) and offspring produced by older parents may experience greater telomere loss during the growth period (B. Heidinger et al., 2016).

Currently there is no synthesis of the published literature summarizing the prevalence and magnitude of parental age at conception effects on offspring TL across studies and species. This is despite the growing interest in the topic of how parental age impacts offspring longevity, especially because of the increasing average age of offspring production in human populations (Gillespie et al., 2013). To address this gap, we conducted a meta-analysis to quantify the magnitude of the effects of paternal and maternal age at offspring conception on offspring TL in vertebrates. We included data from published literature on vertebrates, tested for both parental and offspring sex-specific effects, controlled for phylogeny and standardized effect estimates, allowing for greater accuracy and comparability across studies.

We tested whether there was an association between offspring TL and parental age at offspring conception across vertebrate species. We also tested whether the association varies across species, telomere extraction methods, statistical methods, and both parental and offspring sex. Furthermore, we tested for publication and time-lag biases to assess the validity of any general conclusions drawn from existing studies. We hypothesized that there would be a positive association between paternal age and offspring TL (Eisenberg & Kuzawa, 2018) and a negative association between maternal age and offspring TL (Keefe, 2016).

## Materials and Methods

### Systematic review

We conducted data collection and analyses using R 4.4.1 (R Core Team, 2024), unless otherwise stated. We performed a systematic literature search for publications using *litsearchr* 1.0.0 (Grames et al., 2019) and the Web of Science database up to November 2023 (last publication date October 13, 2023). Following the *litsearchr* guidelines (Grames et al., 2019), a list of 27 highly relevant articles was obtained from a naive search string on the Web of Science (ALL=(“parental age” AND “offspring” AND “telomere length”) AND DT=(Article)). The search terms, which consisted of at least one word that occurred at least three times across these articles, were extracted from article titles, abstracts, and authors’ keywords. The resulting terms were manually selected and grouped to create the final Boolean search string, which was: (ALL=((matern* OR mother* OR patern* OR father* OR parent*) AND (age* OR aging OR senesc* OR old* OR young* Or effect* OR “age at concept*”) AND (transgener* OR juvenil* OR offspr* OR child* OR birth* OR young*) AND (“lans* effect*” OR “telomere *”))). This search returned 871 candidate articles with 3 duplicates which we removed, resulting in a total of 868 articles, which were passed onto the screening process (Figure 1).

**Figure 1.**
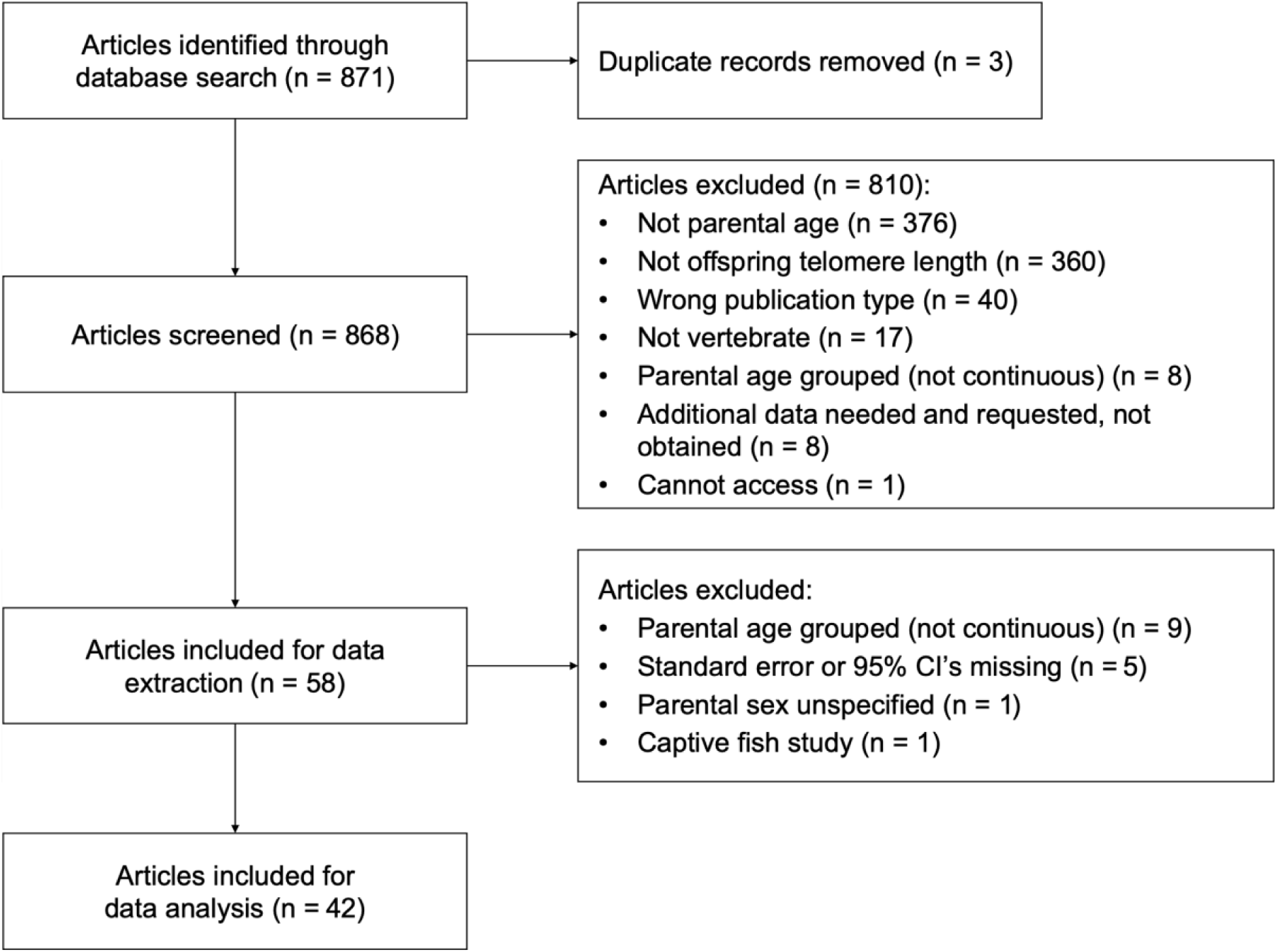
Modified preferred reporting items for systematic reviews and meta-analyses (PRISMA) flow chart showing the screening process used to exclude or retain articles for this meta-analysis (Moher, 2009)

Title and abstract screenings were conducted using the Rayyan web-based application (Ouzzani et al., 2016). Studies passing this initial screening had their full text reviewed to be included in the meta-analysis. Studies were included if they 1) were original research articles; 2) used a vertebrate; 3) used continuous separate paternal or maternal age at conception as explanatory variable (instead of categorical age groups or a measure of combined parental ages); 4) measured offspring telomere length as the response variable; and 5) reported effect sizes with associated standard errors (SE) or 95% confidence intervals (CI), or reported correlation coefficients, or provided raw data (n=1) to reconstruct the statistical model described in the text to calculate the effect size and corresponding error (Figure 1). Initial and full-text screenings were done by two researchers independently, and any conflicts were identified, discussed, and resolved. Fifty eight articles remained after screening.

### Data extraction

For the retained articles, effect size estimates and their corresponding standard errors were extracted or calculated. Slopes were extracted if the article used linear regression as their statistical method. For articles that reported multiple models estimating the same effect (for example, various models controlling for different factors but all estimating the parental age effect on offspring TL), the estimate was extracted from the model identified by the authors as the best-fitting or otherwise the most complex model with the largest number of parameters to account for confounding factors. Correlation coefficients were extracted if the article used correlation as their statistical method. When only 95% confidence intervals were provided, the standard error was calculated using the formula (upper interval limit - estimate)/1.96.

Data were extracted from the main text, supplementary information, relevant figures and tables. For each effect size estimate, the following were recorded: article publication date, authors, title, study ID, estimate ID (a unique number for each estimate if there were multiple estimates made in one study), species, offspring sample size, number of observations, parental sample size (i.e. number of mothers or number of fathers depending on the focal parental sex), offspring sex (“son”, “daughter” or “unspecified” if the article did not distinguish between the offspring sex), parental sex (“father” or “mother”), sperm production seasonality (“seasonal” or “continuous”), environmental conditions (“human” for human studies, “captive” for captive vertebrate populations, or “wild” for wild vertebrate populations), offspring age at measurement (“juvenile” if the measuring occurred before the offspring reached sexual maturity, “adult” for after sexual maturity, or “both” if a mixture of the two other categories), laboratory method (“qPCR”, “TRF Southern blot”, “TRF in gel”, or “WGS”), whether parental environment effects were controlled for in the analysis (e.g. disease, smoking; “yes” or “no”), cell type from which telomeres were extracted (“leukocytes”, or “erythrocytes”, or “other”; “other” included “umbilical cord” from a single study, “sperm” from two studies, and “buccal cells” from two studies), whether the parent’s identity was controlled for (“yes” or “no”), whether the effect size estimate was separated into within- and between-parent effects, and if yes, which component the estimate corresponded to (“within”, or “between”, or “neither” if not separated). In addition, we recorded additional information needed to convert the effect size estimates for further analysis: statistical methods (“regression” or “correlation”), number of parameters and degrees of freedom used in the study’s model. As a control, two researchers independently selected a random subset of 10 articles and extracted these values. This random subset had 33 discrepancies out of 1062 (3%) values which were subsequently reviewed and resolved.

In 9 cases where the article mentioned testing the parental age effect on offspring TL but did not report the corresponding effect sizes and standard errors, authors of those studies were contacted to obtain the missing data. Studies that did not provide the requested information (8 studies) were excluded from further analysis. We removed a study conducted in salmon fry under captive conditions because it was the only study in our dataset that did not account for the telomere dynamics under natural conditions (McLennan et al., 2018). The consequential dataset contained 5297 values from 42 studies to record (Figure 1).

### Effect size calculation

To allow comparison between studies using regression and correlation methods, slope estimates were converted to correlation coefficients using the formula

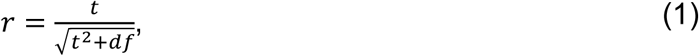

where *t* is the t-value of the linear regression and *df* is the degree of freedom (Nakagawa & Cuthill, 2007; Remot et al., 2021). To correct potentially skewed sampling distribution, the reported and calculated correlation coefficients were transformed to population correlations, following the formula

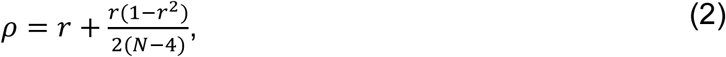

where *r* is the correlation coefficient and N is the sample size used to obtain the correlation coefficient (Gnambs, 2023). These were subsequently transformed to Fisher’s *z* (*Zr*) (Nakagawa & Cuthill, 2007; Remot et al., 2021) using “ZCOR” argument in the escalc function from *metafor* 4.6-0 (Viechtbauer, 2010), which follows the formula

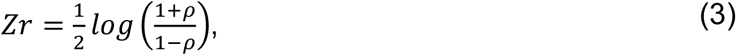

where *ρ* is the population correlation and represents the adjusted correlation coefficient instead of *r*. The resulting Zr was close to being normally distributed (mean = 0.06, SD = 0.17, range= -0.72 to 0.75; Figure 2). Sampling variances for each Zr were computed using the same “ZCOR” argument in the *escalc* function, which follows the formula

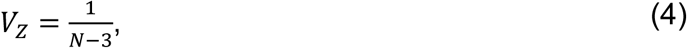

where N is the sample size used to obtain the correlation coefficient (Nakagawa et al., 2021). To account for phylogenetic relationships in this meta-analysis, a phylogenetic tree (Figure 3) was constructed with species data from the Open Tree of Life using *rotl* 3.1.0 (Michonneau et al., 2016) and a phylogenetic correlation matrix using *ape* 5.8 (Paradis & Schliep, 2018), and visualized using *ggtree* 3.12.0 (Yu et al., 2017).

**Figure 2.**
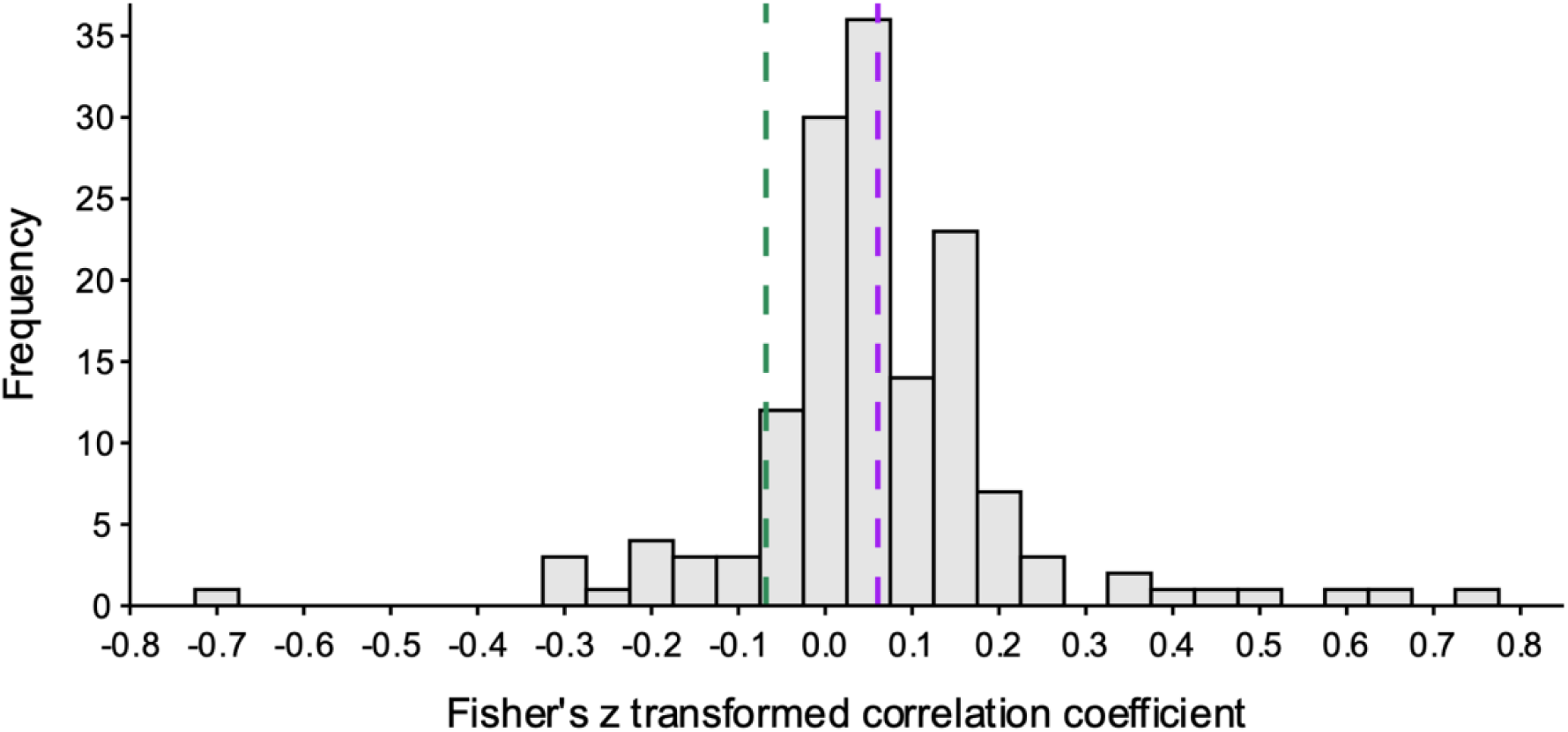
Histogram displaying the frequency distribution of Fisher’s z transformed correlation coefficients (Zr) from the dataset (n = 148). Purple dashed line indicates unadjusted Zr, green dashed line indicates the adjusted Zr for study, estimate, and phylogenetic effects.

**Figure 3.**
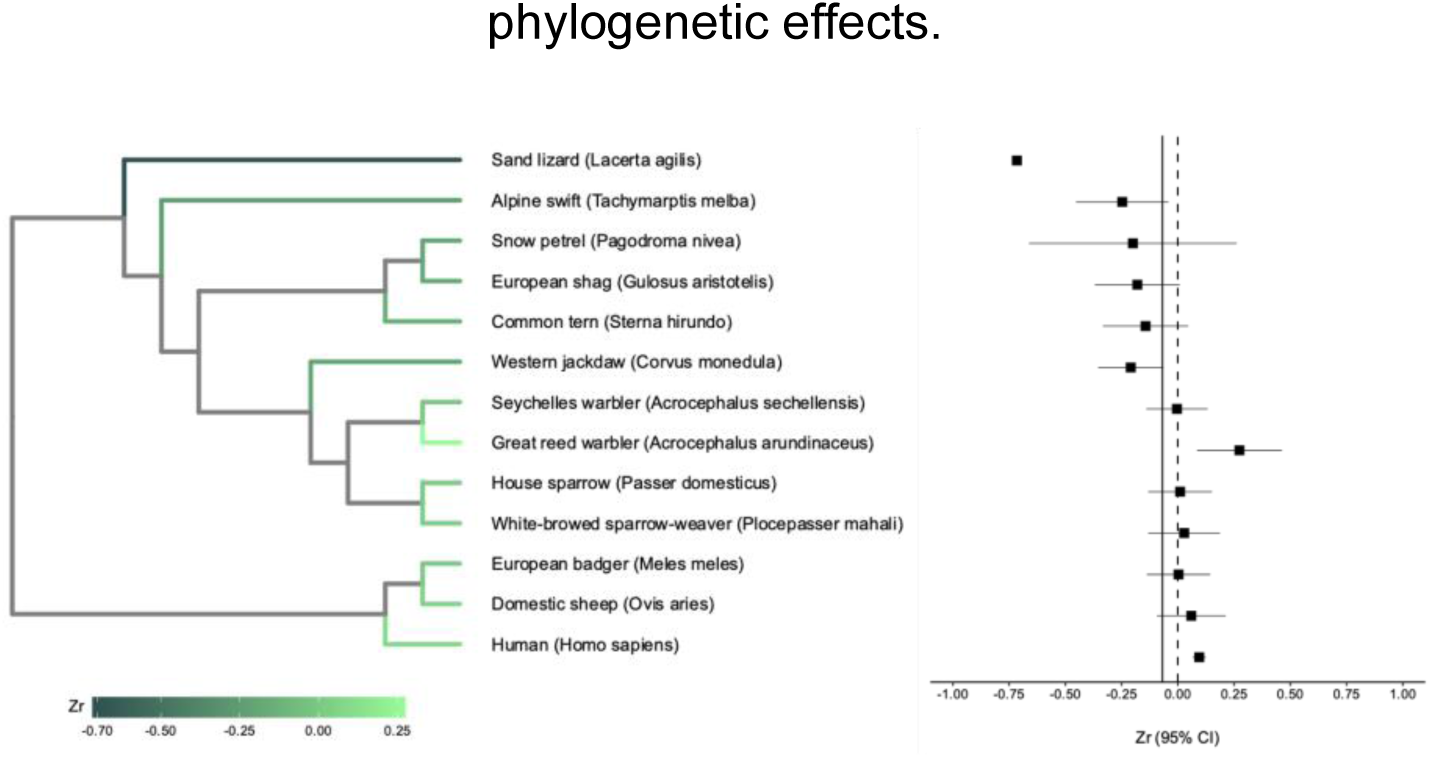
Phylogenetic tree (left) and forest plot (right) of the 13 vertebrate species included in this meta-analysis with corresponding adjusted mean Zr. The phylogenetic tree green color represents species-level predicted Zr values, with darker shades indicating lower effect sizes, and lighter shades indicating higher effect sizes. In the forest plot, each dot represents the species-level predicted effect size (Zr) estimated from the meta-regression model, with horizontal lines indicating the respective 95% confidence intervals (CI). The dotted vertical line corresponds to zero and the solid vertical line shows the overall mean-adjusted Zr across species.

### Statistical analysis

The final dataset contained 148 effect sizes from 42 studies and 13 species (Figure 4 aggregated per study; Figure S1 non-aggregated). Studies conducted in humans accounted for 30 of the 42 studies, with the remaining 12 species appearing in one study each. Considering the data imbalance and the fact that non-human studies often revealed patterns inconsistent with findings in human studies, we divided the dataset into human (n = 99 effect sizes) and non-human subsets (n = 49 effect sizes) and modeled them separately to avoid bias toward human studies.

**Figure 4.**
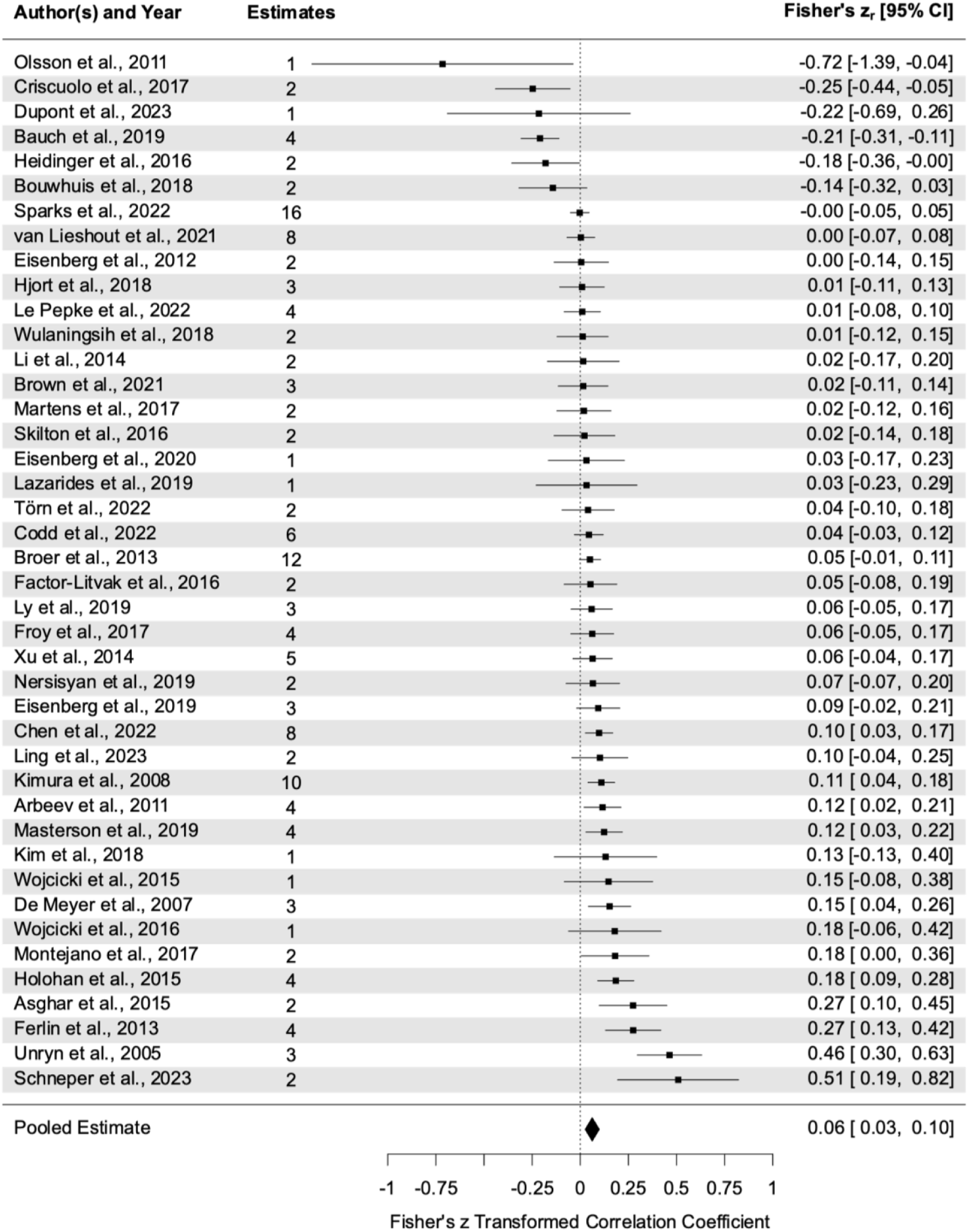
Forest plot of the studies included in the meta-analysis, showing the aggregated effect sizes per study. Each dot represents the cumulative effect size from the study, with horizontal lines indicating the respective 95% confidence intervals (95% CI) and the dotted vertical line indicating zero. The number of effect size estimates for each study is shown in the corresponding column.

The meta-analysis was conducted using *metafor* 4.6 (Viechtbauer, 2010). First, global parental-age effects were quantified using intercept-only models (model 1.1 for the human subset, model 1.2 for the non-human subset), where Zr was fitted as the response variable and the associated sampling variance was used as the weight for each coefficient. To account for non-independence among effect sizes, the following random factors were added to models (Nakagawa & Santos, 2012; Viechtbauer, 2010): study ID, to estimate study-specific effects 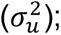 estimate ID, to test for within-study between-estimate variation as the number of effect sizes did not equal the number of studies included in the model 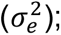 phylogeny weighted by a phylogenetic similarity matrix to test for phylogenetic variation 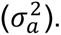 Phylogeny was not included in model 1.1 because there was only one species present. Species is not included in the models because there was either only one species present in the data (model 1.1) or species explained the same variance as study ID (model 1.2).

The typical sampling-error variance (within-study variance, 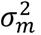), the total model variance 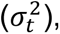 and total estimated model heterogeneity (*I*^2^) were calculated following (Nakagawa & Santos, 2012) with the following formulae:

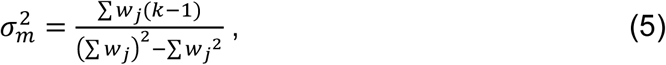

where *w*_*j*_ is the inverse of sampling variance of *j^th^* size and *k* is the number of studies;

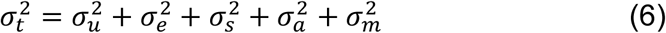

and

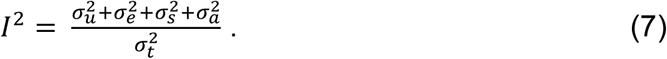

For each random effect the proportion of variance it explains was calculated by

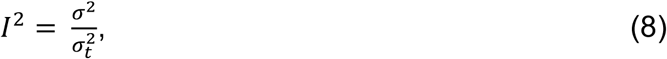

where *σ*^2^ is the respective variance component.

Model 1.1 and 1.2 were then expanded into multilevel random-effects models (model 2.1 and 2.2, respectively) using maximum likelihood (ML) estimation to quantify the effects of the relevant biological and methodological factors. The following moderators were included as fixed effects: offspring sex, parental sex, offspring age, laboratory method, sperm production seasonality, cell from which telomeres were extracted, whether within- or between-parent effects were separated (to account for selective (dis)appearance, (van de Pol & Wright, 2009)), whether parental environment effects were controlled for as they may bias the parental age effect (Lulkiewicz et al., 2020), and whether the parent’s identity was controlled for. Model multicollinearity was assessed using the *vif* function from *metafor* (Viechtbauer, 2010).

As parental effects could depend on the combination of parent and offspring sexes, an interaction of parental sex with offspring sex was added to the models. Because immunosenescence in leukocyte types leads to an increase in the proportion of granulocytes which have longer TL, an interaction between offspring age and telomere cell type was added (Peters et al., 2019; Rufer et al., 1999). The random effect structure stayed the same as in the intercept-only model. There was no significant interaction between cell type and offspring age, and this interaction was removed from the models. In the non-human vertebrate subset, the interaction between parental and offspring sex was not included because of the lack of studies specifically testing parental age effects on daughters. An omnibus test of all model coefficients excluding the intercept was run to test whether the variance was sufficiently explained by the moderators (Viechtbauer, 2010). For laboratory methods and the interaction between parental and offspring sexes, posthoc tests were conducted using *emmeans* 1.10.7 to enable significance tests among all levels (Lenth, 2025). The percentage of heterogeneity explained by each moderator was calculated following (Nakagawa & Schielzeth, 2013) as marginal R^2^ using *orchaRd* 2.0 (Nakagawa et al., 2023).

### Publication and time-lag biases

To test for publication bias in the human and non-human vertebrate studies, contour-enhanced funnel plots made with *metafor* (Viechtbauer, 2010) were visually assessed. The plots display precision (inverse of the standard error) on the y-axis against the residuals of the Zr derived from the full meta-regression model. Plotting residuals from the meta-regression removes some of the heterogeneity due to confounding factors like study design or phylogenetic relationships, which could otherwise cause misleading asymmetry (Nakagawa et al., 2021). A symmetric distribution of data points around the mean effect size suggests the absence of publication bias (Nakagawa et al., 2021).

Publication bias was also assessed using a modified Egger’s regression test (Egger et al., 1997) on a uni-moderator model using sampling variances of the effect sizes as a predictor (Nakagawa et al., 2021). To test for potential time-lag bias, the models were rerun with a mean-centered year of publication as a moderator (Nakagawa et al., 2021). Included random effects were the same as in the main models (study ID and estimate ID in the human model, plus phylogeny in the non-human model) to account for the non-independence of estimates across studies (Nakagawa et al., 2021).

## Results

### Intercept only models (models 1.1 & 1.2)

Model 1.1 detected a statistically significant and positive parental age effect on offspring TL in the human subset, which yielded an estimated Zr of 0.095 (95% CI = 0.067, 0.124), after adjusting for study and estimate effects (Table 1a). The I^2^ value was 98.6%, indicating substantial heterogeneity across correlation coefficients, mainly due to between-study effects (65%, Table 1a). No significant parental age effect on offspring TL was found in the non-human subset (model 1.2), which yielded an estimate of Zr of -0.087 (95% CI = -0.267, 0.094; Table 1b). The I^2^ was 99.0%, indicating considerable heterogeneity across correlation coefficients, which was mainly due to phylogeny (75%) and study (19%) effects (Table 1b).

**Table 1.**
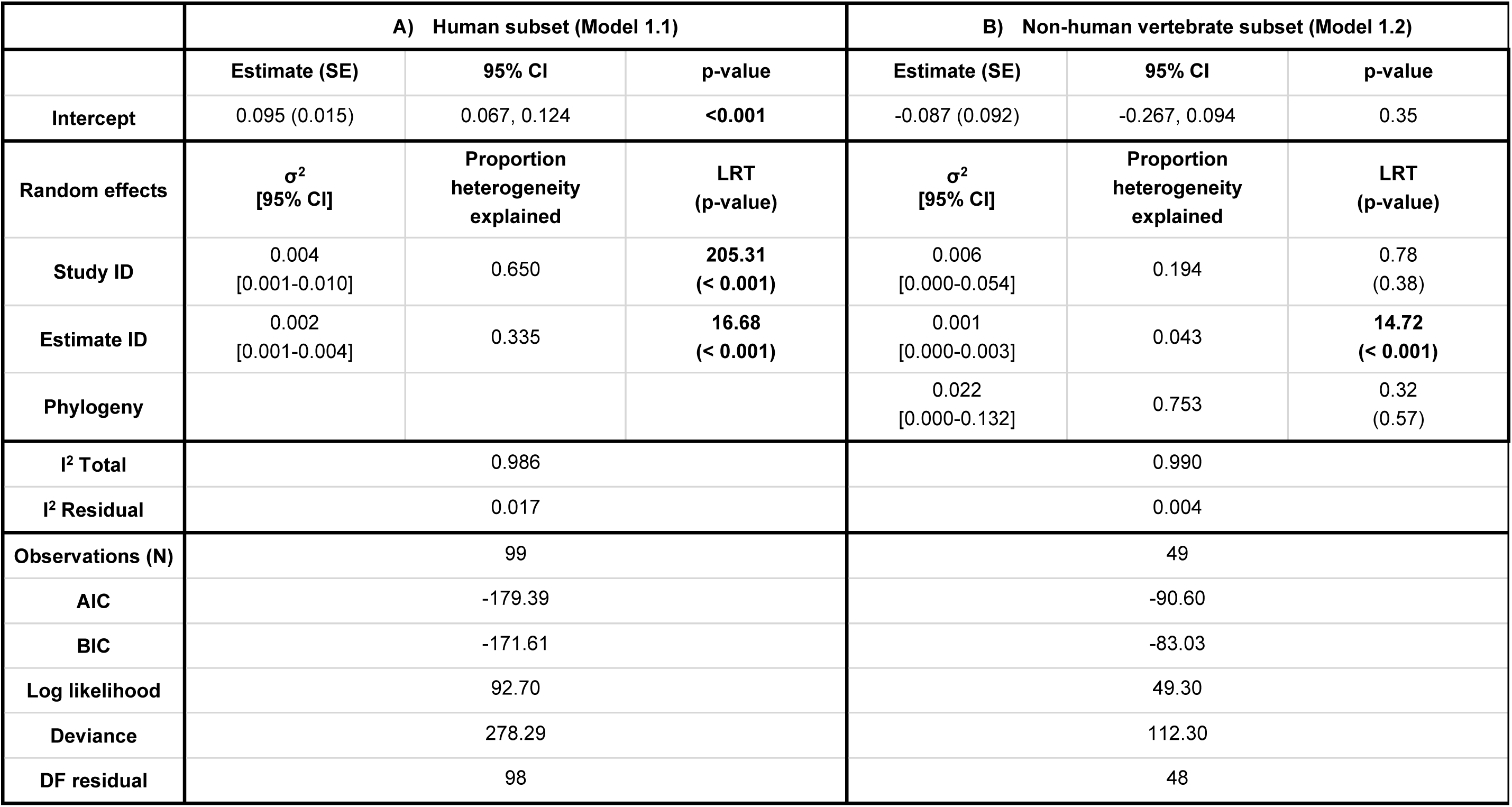
Variance among studies, estimates, species and phylogeny from the meta-analytic intercept-only models of parental age at conception effects on offspring telomere length in A) humans (model 1.1) and B) 12 non-human vertebrates (model 1.2). Significant intercept and random effects are in bold. SE = standard error, 95% CI = confidence interval, DF = degrees of freedom.

|  | A) Human subset (Model 1.1) |  |  | B) Non-human vertebrate subset (Model 1.2) |  |  |
| --- | --- | --- | --- | --- | --- | --- |
|  | Estimate (SE) | 95% CI | p-value | Estimate (SE) | 95% CI | p-value |
| Intercept | 0.095 (0.015) | 0.067, 0.124 | **<0.001** | -0.087 (0.092) | -0.267, 0.094 | 0.35 |
| Random effects | $\sigma^2$<br>[95% CI] | Proportion<br>heterogeneity<br>explained | LRT<br>(p-value) | $\sigma^2$<br>[95% CI] | Proportion<br>heterogeneity<br>explained | LRT<br>(p-value) |
| Study ID | 0.004<br>[0.001-0.010] | 0.650 | **205.31**<br>(**< 0.001**) | 0.006<br>[0.000-0.054] | 0.194 | 0.78<br>(0.38) |
| Estimate ID | 0.002<br>[0.001-0.004] | 0.335 | **16.68**<br>(**< 0.001**) | 0.001<br>[0.000-0.003] | 0.043 | **14.72**<br>(**< 0.001**) |
| Phylogeny |  |  |  | 0.022<br>[0.000-0.132] | 0.753 | 0.32<br>(0.57) |
| I<sup>2</sup> Total | 0.986 |  |  | 0.990 |  |  |
| I<sup>2</sup> Residual | 0.017 |  |  | 0.004 |  |  |
| Observations (N) | 99 |  |  | 49 |  |  |
| AIC | -179.39 |  |  | -90.60 |  |  |
| BIC | -171.61 |  |  | -83.03 |  |  |
| Log likelihood | 92.70 |  |  | 49.30 |  |  |
| Deviance | 278.29 |  |  | 112.30 |  |  |
| DF residual | 98 |  |  | 48 |  |  |

### Full meta-regression models (models 2.1 & 2.2)

Sample sizes and distributions of the estimates for each level of the moderators are shown in Figures 5 & S2 (human subset) and Figures 6 & S3 (non-human vertebrate subset). The I^2^ for model 2.1 was 95% (Table 2a), where most of the heterogeneity was explained by estimate ID (75%). The omnibus test of moderators was highly significant (QM_df = 13_ = 74.5, p-value < 0.001), suggesting that the included moderators explain significant variability in the model (Viechtbauer, 2010).

**Figure 5.**
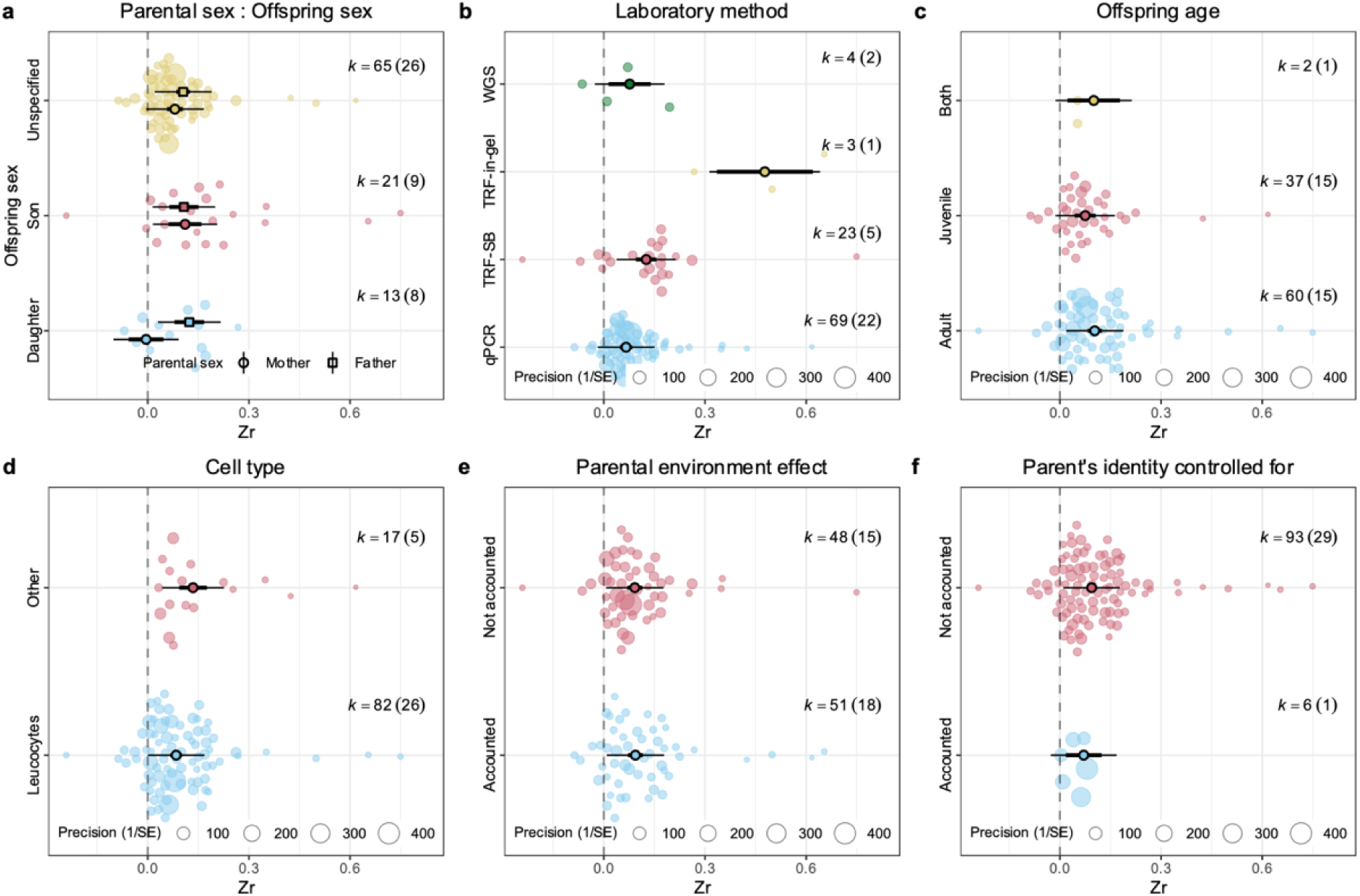
Orchard plots showing the estimated effect sizes across (a) parental and offspring sexes (reference level = Mother and Daughter), (b) laboratory method (reference level = qPCR), (c) offspring age at measurement (reference level = Adult), (d)cell type from which the telomeres were extracted (reference level = Leukocytes), (e) whether parental environment effects were controlled for (reference level = No), and (f) whether the parent’s identity was controlled for (reference level = No) in the human data subset. In each plot, the solid dot indicates the estimated mean effect size predicted by the meta-regression model, the thick whiskers indicate the 95% confidence intervals, the thin whiskers indicate the 95% prediction intervals (the range in which the point estimate of 95% of future studies will fall), and translucent dots indicate the distribution of the raw effect sizes. The size of the translucent dots indicate precision of the effect size. K indicates the number of effect sizes, and numbers in parentheses indicate the number of studies.

**Figure 6.**
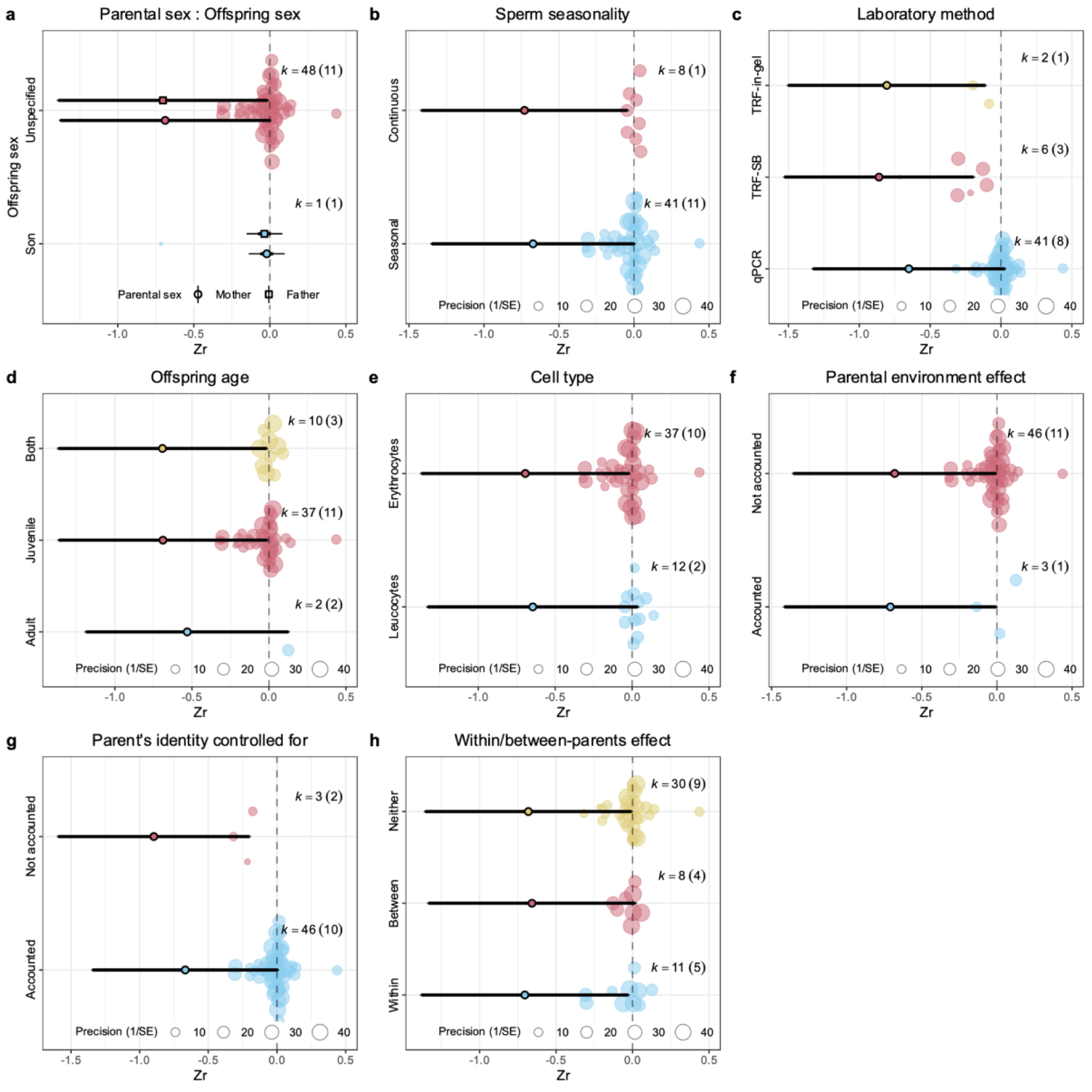
Orchard plots showing the estimated effect sizes across (a) parental and offspring sexes (reference level = Mother and Unspecified-sex offspring; note, there were no studies on only daughters and there was no significant interaction between parent and offspring sex), (b) sperm production seasonality (reference level = continuous), (c) laboratory method (reference level = qPCR; SB = southern blot), (d) offspring age at measurement (reference level = Adult), (e) cell type from which the telomeres were extracted (reference level = Leukocytes), (f) whether parental environment effects were controlled for (reference level = No), (g) whether the parent’s identity was controlled for (reference level = No), and (h) whether within-/between-parent effects were accounted for (reference level = Neither) in the non-human vertebrate data subset. In each plot, the solid dot indicates the estimated mean effect size predicted by the meta-regression model, the thick whiskers indicate the 95% confidence intervals, the thin whiskers indicate the 95% prediction intervals (the range in which the point estimate of 95% of future studies will fall), and translucent dots indicate the distribution of the raw effect sizes. The size of the translucent dots indicate precision of the effect size. K indicates the number of effect sizes, and numbers in parentheses indicate the number of studies.

**Table 2.** Moderator estimates from the meta-regression models of parental age effect on TL in: A) humans (model 2.1 - human subset) and B) 12 non-human vertebrates (model 2.2 - non-human vertebrate subset). Both models account for study and estimate, and for B) phylogenetic non-independence. Non-significant interactions were removed. Parentheses in the fixed effects subheadings indicate the reference level; p-values <0.05 are in bold and <0.1 are in italics. SE = standard error, 95% CI = confidence interval, DF = degrees of freedom. TRF = terminal restriction fragments, WGS = whole genome sequencing.

|  | A) human subset (Model 2.1) |  |  | B) on-human vertebrate subset (model 2.2 ) |  |  |
| --- | --- | --- | --- | --- | --- | --- |
| Fixed effects | Estimate (SE) | [95% CI] | p-value | Estimate (SE) | [95% CI] | p-value |
| intercept | 0.029 (0.031) | [-0.090, 0.031] | 0.56 | -0.059 (0.131) | [-0.316, 0.197] | 0.65 |
| Parental sex (Mother) |  |  |  |  |  |  |
| Father | 0.128 (0.031) | [0.067, 0.189] | **< 0.001** | -0.015 (0.014) | [-0.042, 0.012] | 0.28 |
| Offspring sex (Female for 2.1, unspecified-sex for 2.2) |  |  |  |  |  |  |
| Unspecified-sex offspring | 0.085 (0.030) | [0.027, 0.144] | **0.004** |  |  |  |
| Male offspring | 0.116 (0.033) | [0.051, 0.181] | **< 0.001** | -0.668 (0.353) | [-0.023, 1.359] | *0.06* |
| Sperm production seasonality (Continuous) |  |  |  |  |  |  |
| Seasonal |  |  |  | 0.057 (0.082) | [-0.103, 0.217] | 0.48 |
| Parental environment controlled for (No) |  |  |  |  |  |  |
| Present | 0.002 (0.018) | [-0.032, 0.036] | 0.91 | -0.029 (0.082) | [-0.189, 0.131] | 0.72 |
| Parent's identity controlled for (No) |  |  |  |  |  |  |
| Yes | -0.024 (0.028) | [-0.079, 0.032] | 0.40 | 0.228 (0.090) | [0.053, 0.404] | **0.01** |
| Within-/ between-parent effects (Neither) |  |  |  |  |  |  |
| Between |  |  |  | 0.023 (0.023) | [-0.019, 0.065] | 0.28 |
| Within |  |  |  | -0.023 (0.022) | [-0.065, 0.019] | 0.29 |
| Offspring age at measurement (Adult) |  |  |  |  |  |  |
| Both | -0.003 (0.039) | [-0.079, 0.074] | 0.94 | -0.161 (0.083) | [-0.323, 0.002] | *0.05* |
| Juvenile | -0.028 (0.022) | [-0.071, 0.015] | 0.20 | -0.158 (0.081) | [-0.316, 0.000] | *0.05* |

| Laboratory method (qPCR) |  |  |  |  |  |  |
| --- | --- | --- | --- | --- | --- | --- |
| TRF in-gel | 0.411 (0.073) | [0.267, 0.555] | < 0.001 | -0.156 (0.087) | [-0.327, 0.015] | 0.07 |
| TRF Southern blot | 0.060 (0.019) | [0.022, 0.098] | 0.002 | -0.210 (0.062) | [-0.332, -0.089] | < 0.001 |
| WGS | 0.011 (0.033) | [-0.054, 0.076] | 0.75 |  |  |  |
| Telomere cell type (Leukocytes) |  |  |  |  |  |  |
| Erythrocytes |  |  |  | -0.048 (0.069) | [-0.184, 0.087] | 0.48 |
| Other | 0.050 (0.024) | [0.003, 0.097] | 0.04 |  |  |  |
| Interaction parental sex:offspring sex (Mother : Daughter) |  |  |  |  |  |  |
| Father : Son | -0.132 (0.043) | [-0.216, -0.048] | 0.002 |  |  |  |
| Father : Unspecified-sex offspring | -0.104 (0.034) | [-0.170, -0.038] | 0.002 |  |  |  |
| Random effects | Estimate ($\sigma^2$) | 95% CI | Proportion heterogeneity explained | Estimate ($\sigma^2$) | 95% CI | Proportion heterogeneity explained |
| Study ID | 0.000 | 0.000, 0.002 | 0.198 | 0.003 | 0.000, 0.018 | 0.721 |
| Estimate ID | 0.001 | 0.001, 0.003 | 0.753 | 0.001 | 0.000, 0.003 | 0.193 |
| Phylogeny |  |  |  | 0 | 0.000, 0.011 | < 0.001 |
| I<sup>2</sup> total | 0.950 |  |  | 0.914 |  |  |
| I<sup>2</sup> residual | 0.050 |  |  | 0.034 |  |  |
| Observations (N) | 99 |  |  | 49 |  |  |
| AIC | -204.85 |  |  | -89.16 |  |  |
| BIC | -163.33 |  |  | -58.89 |  |  |
| Log likelihood | 118.43 |  |  | 60.58 |  |  |
| Deviance | 226.83 |  |  | 89.74 |  |  |
| DF residual | 85 |  |  | 36 |  |  |

Laboratory method, parental sex and offspring sex and their interaction, and cell type were significant in model 2.1 (Table 2a). Laboratory method explained 62% of the heterogeneity (Table 3). Studies using TRF in-gel (Zr = 0.41, Table 2a) had more positive estimates compared to studies using the other three methods, and TRF southern blot (Zr = 0.06, Table 2a) had more positive estimates compared to studies using qPCR (Figure 7, Table S2). The interaction between parental and offspring sex explained 14% of the heterogeneity (Table 3). Parental age effects were more positive for all parent-offspring sex combinations compared to mothers-daughters (Figure 5a), and the significance was tested by post-hoc tests (Table S3). Cell type explained 4% of the heterogeneity. Studies using cells other than blood cells to extract telomeres had more positive estimates (Table 2a). Whether the parental environment effects were controlled for, whether the parent’s identity was controlled for and offspring age at measurement did not significantly alter the estimated correlation between parental age at conception and offspring TL (Table 2a).

**Figure 7.**
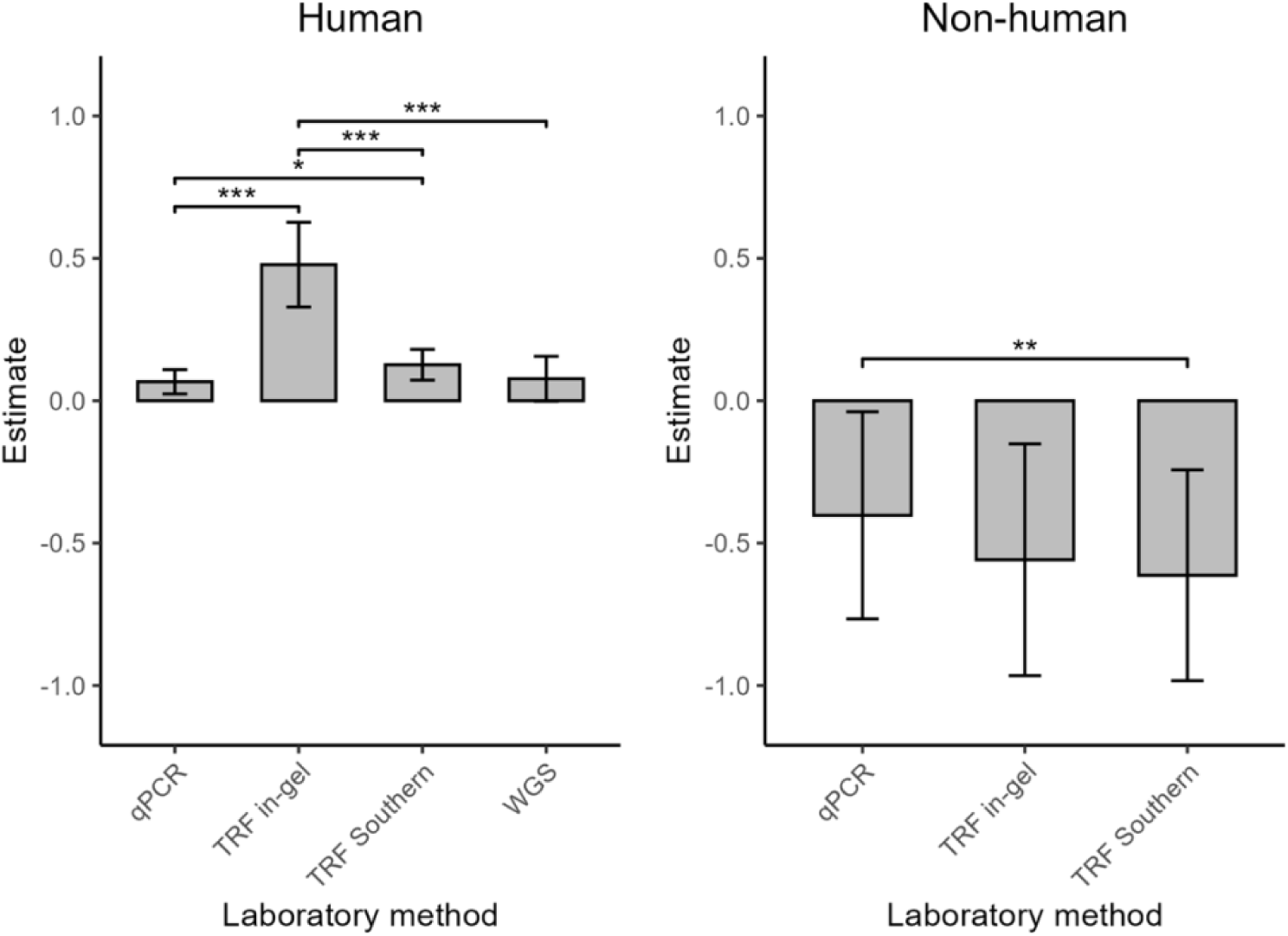
Parental age at conception effects on offspring telomere length estimated in studies using different laboratory methods. Error bars indicate 95% confidence intervals. *0.05 > p > 0.01, **0.01 > p > 0.001, ***p < 0.001 from post-hoc analyses (Table S2). TRF = terminal restriction fragments, WGS = whole genome sequencing.

**Table 3.** R^2^ marginal per each moderator in full meta-regression models.

|  | Model 2.1 (human) | Model 2.2 (non-human) |
| --- | --- | --- |
| Parental sex : offspring sex | 0.1423 | — |
| Offspring sex | 0.0466 | 0.3456 |
| Parental sex | 0.0511 | 0.0029 |
| Sperm production seasonality | — | 0.0011 |
| Parental environment controlled for | 0.0011 | 0.0048 |
| Parent's identity controlled for | 0.0251 | 0.1523 |
| Within-/ between-parent effects | — | 0.0069 |
| Offspring age at measurement | 0.0838 | 0.0231 |
| Laboratory method | 0.6186 | 0.3980 |
| Telomere cell type | 0.0357 | 0.1775 |

The I^2^ for model 2.2 was 91% (Table 2b), where most of the heterogeneity was explained by between-study variance (72%). The omnibus test of moderators was highly significant (QM_df = 13_ = 114.85, p-value < 0.001). Significant moderators for model 2.2 were laboratory methods and parent’s identity controlled for (Table 2b). The laboratory method explained 40% of the heterogeneity (Table 3). Studies using TRF southern blot had more negative estimates compared to studies using qPCR (Table 2b), which was the opposite of model 2.1 (the human subset). Studies using TRF in-gel were marginally non-significant compared to studies using qPCR and not significant compared to TRF Southern blot studies (Figure 7, Table S2). Whether the parent’s identity was controlled for explained 15% of the heterogeneity (Table 3). Estimates were less negative when the parent’s identity was controlled for (Zr = 0.23, Table 2b) than when it was not. Parental sex, whether the parental environment was controlled for, sperm seasonality, whether the effect was between- or within-parent and cell type did not affect the parental age effect estimates, and both offspring sex and offspring age when TL was measured were marginally non-significant (Table 2b). For full model output see Table S1.

### Publication and time-lag biases

Visually, there was asymmetry in the funnel plots, with more published studies present on the right side, suggesting a lack of published studies with effect sizes smaller than the mean residual Zr (Figure 8). Funnel asymmetry was also detected by weighted Egger’s regression using sampling variance as one of the predictors to test for outcome reporting bias, which was found in the human subset (β = 10.89, p-value < 0.01) but not in the non-human vertebrate subset (β = 5.55, p-value = 0.24). Time-lag bias was detected in the human subset (β = 0.005, p-value = 0.05) but not in the non-human vertebrate subset (β = -0.002, p-value = 0.79).

**Figure 8.**
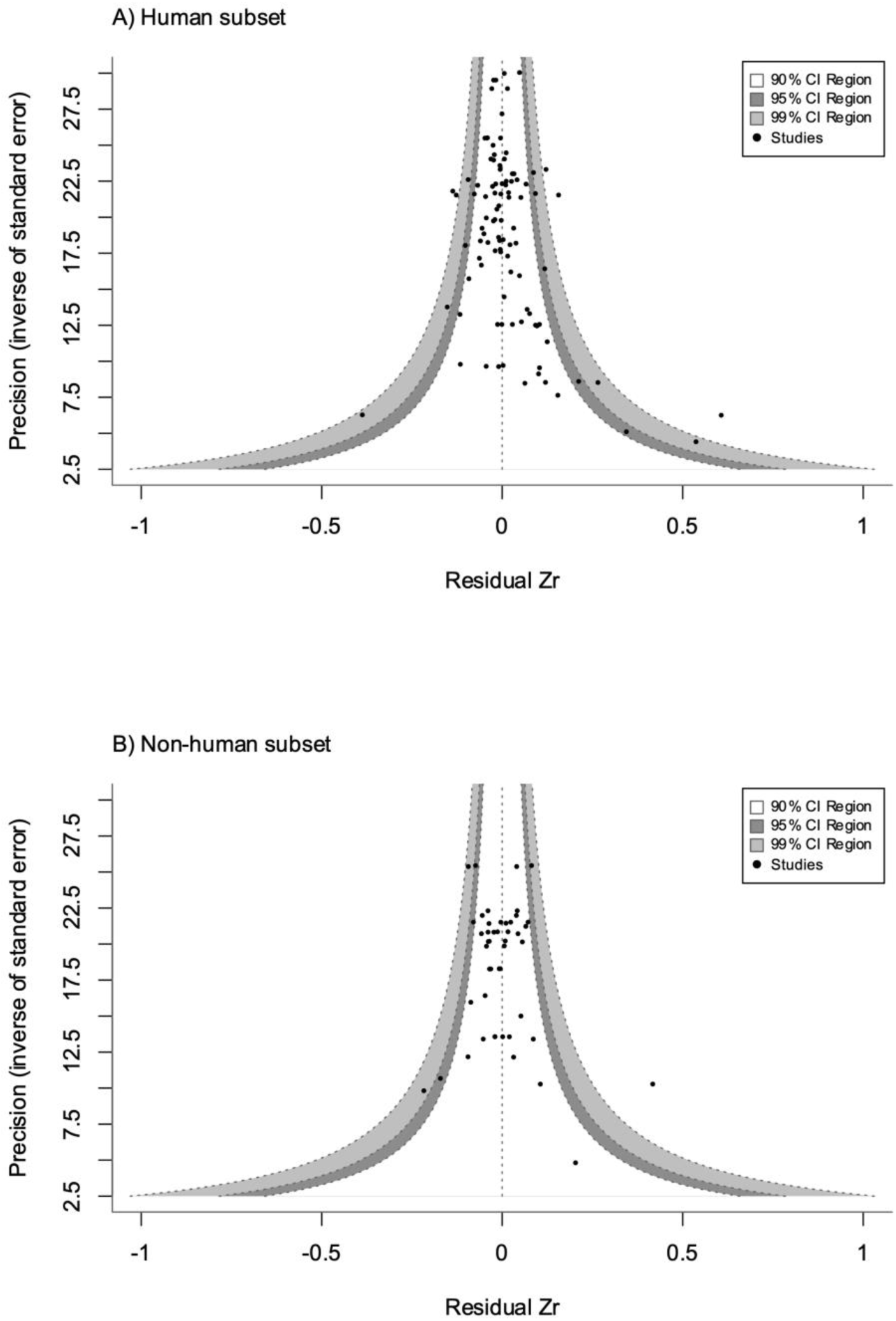
Contour-enhanced funnel plot for (a) human subset and (b) non-human vertebrate subset, with the precision (the inverse of the standard error) on the y-axis against the residuals of the Fisher’s z-transformed correlation coefficients from the full meta-regression model. Dashed vertical line indicates the mean Zr. Shaded gray areas indicate confidence intervals. Dots represent studies used in this meta-analysis.

## Discussion

We performed meta-analyses to assess the overall strength of association and variation sources between parental age at conception and offspring TL in humans and non-human vertebrates in the published literature. We found a significant positive (humans) and no (non-human vertebrates) parental age effect on offspring TL. Laboratory method explained the most heterogeneity in both analyses (62%, humans; 40%, non-human vertebrates) but the direction of the effects differed. Parental sex, offspring sex, and their interaction, and cell type affected the parental age effect estimates in humans but not in non-humans. Whether the parent’s identity was controlled for in the study affected the parental age effect estimates in non-humans but not in humans. This highlights moderators that are important to account for in future studies, and the different effects that may have across taxa.

### Source of heterogeneity

Substantial heterogeneity was observed in the estimates of parental age effect on offspring TL. In the human subset (model 1.1), the among-study variation accounted for most of the heterogeneity (65%, Table 1a); in the non-human vertebrate subset (model 1.2), phylogeny was the major contributor (76%, Table 1b). This suggests that in human studies, most of the heterogeneity was attributed to study-specific factors, such as differences in laboratory methods and similarities within the study populations. In non-human vertebrate studies, the high phylogenetic signal, which was also reflected by the visual trend in the colour of the phylogenetic tree (Figure 3), suggests that shared evolutionary history as well as similar ecological niches and life-history strategies may account for much of the consistency in parental age effects on offspring TL in closely related species. However, phylogeny was not significant and therefore this result should be interpreted with caution. This may be partly due to the limited and unbalanced taxonomic coverage of the non-human vertebrate subset, which contained 9 birds, two mammals and one lizard, highlighting a taxa imbalance that needs addressing in the literature. In particular, future studies on amphibians and fish are required. Additionally, since there was only one study for each species in the non-human vertebrate subset, we could not distinguish between the study effect and species effect. To accurately determine phylogenetic patterns in the effects of parental age on offspring TL, further research is needed in non-human vertebrate taxa.

### Methodological factors affecting parental age effect estimates

Effect size estimates differed by telomere measurement methods in both human and non-human studies. In human studies (model 2.1), the laboratory method moderator explained a large proportion of the variation: TRF with in-gel hybridization has a large positive effect on Zr (0.411, Table 2a), and a large R^2^ (0.619, Table 3). TRF methods, especially TRF with in-gel hybridization, introduce less noise than qPCR (Kärkkäinen et al., 2021; Nussey et al., 2014), and therefore TRF methods are more likely than qPCR to detect an effect if there is one. In non-human studies, the direction of the effects differed with TRF Southern blot having more negative estimates than qPCR, and the estimates derived from TRF in-gel did not significantly differ from those derived from qPCR (although there was a marginal non-significant trend) or TRF Southern blot (Table 2a, Figure 7). This could be due to the small sample size of estimates using TRF in-gel (n = 2), thus the effect of TRF in-gel might be confounded with study- and species-specific effects. Our results show that TRF-based methods better detect associations between parental age at conception and offspring TL, and are recommended for future studies where feasible; otherwise, carefully accounting for the laboratory method used is essential.

We only found five studies out of 42 that separated within-parent from between-parent effects, and none of them were in humans. Therefore, the overall positive effect of parental age at conception in humans is likely to be due to among-individual difference, i.e. parents who gave birth at older ages were of better quality, in both biological and social economic aspects (Boivin et al., 2009; Sanghvi et al., 2024). The offspring might inherit advantageous genes or benefit from an abundance of resources, both of which contribute to better somatic status, associated with longer telomeres (Eisenberg & Kuzawa, 2018; Monaghan & Metcalfe, 2019; Sanghvi et al., 2024). The lack of studies separating within- and between-parent effects restricted our ability to assess the influence of parental age on offspring TL on a within-parent level. While within-parent effects assess changes in offspring TL related to age within the same parent over time, between-parent effects compare offspring TL outcomes across parents of varying ages, and so accounting for these distinctions is necessary to prevent potential biases introduced by among-individual difference when younger or older parents might be overrepresented in specific studies.

In human studies, cell type from which the telomeres were extracted also affected the conclusion. “Other” cell types (umbilical cord, buccal cells and sperm) had a more positive parental age effect on offspring telomere length than leukocytes. Among human organs, TL is shortest in blood and longest in the testis, and the negative association between TL and age is strongest for cell types with shorter average TL (Demanelis et al., 2020). This may explain why leukocytes had smaller estimates than other cell types in our study, because the effect of parental age at conception was reduced due to the short leukocytes TL and the strong effect of offspring age. This result should also be interpreted with caution, given that we only had 5 out of 31 studies using other cell types. In non-human vertebrates, there was no difference between erythrocytes (9 birds and one lizard) and leukocytes (two mammal species), controlling for phylogeny, but again this analysis had limited statistical power and tissue representation. All of the studies used blood-derived TL. While it is a proxy for TL in most tissues, blood-derived TL may not fully capture patterns present in other somatic tissues such as proliferative tissues where telomerase is active (Haussmann et al., 2007; Li, 2012). Future research could benefit from incorporating multiple cell types within the same study design to better assess and potentially reduce this cell type-related bias.

The size of the parental age effect on offspring TL was dependent on the interaction between parental sex and offspring sex in humans, but not in the non-human subset. In humans, parental age effects on offspring TL were more positive for all combinations of parental sex and offspring sex, compared to mother-daughters (Figure 5a, Table S3). This can be discussed from two perspectives: (A) why paternal age effect is more positive than maternal age effect in daughters; and (B) why daughters and sons respond differently to maternal age. Paternal and maternal age effects could differ in their directions, which leads to (A). On one hand, positive paternal age effects could exist due to selective survival or replication of sperm precursor cells (spermatogonia) with longer telomeres as men age, which might lead to a shift in the distribution of TLs within the sperm pool (Eisenberg, 2010; Eisenberg & Kuzawa, 2018). On the other hand, a negative maternal age effect could occur at the same time, which could be explained by oocytes ovulated later in the life cycle being more likely to have more mitotic divisions than those ovulated earlier (Eisenberg & Kuzawa, 2018). However, the difference in the directions of paternal and maternal age effects does not explain (B). Sons and daughters could respond to maternal age differently because of their different environmental sensitivity during development. Advanced maternal age can negatively affect uterine environment (Marti-Garcia et al., 2024; Nelson et al., 2013) and alter milk macronutrient (Lubetzky et al., 2015) and hormone components (Gila-Díaz et al., 2020). While some studies found female fetuses were more sensitive to these prenatal and early postnatal environmental conditions than male fetuses (Vickers et al., 2011), other studies found the opposite (Stinson, 1985). Causes of the absence of the positive parental age effect in the mother-daughter combination remain to be explored.

The insufficient data for offspring sex in non-human vertebrates (Figure S3b) limited our ability to assess the influence of offspring sex and its interaction with parental sex on parental age effect estimates. The sex-specific effect is likely to also exist in non-human vertebrates, because parental care is influenced by parental age, and daughters and sons can differ in their sensitivity to environmental conditions during the growth period (B. J. Heidinger & Young, 2020; Lindström, 1999; Wilkin & Sheldon, 2009). We recommend that future studies take offspring sex into account when investigating parental age effects on offspring TL. Parental age effects through specific parent-offspring sex combinations have been found in other phenotypes such as lifetime reproductive success(Bouwhuis et al., 2015; Kroeger et al., 2020; Schroeder et al., 2015) and longevity (Gavrilov, 1997), although the patterns observed differ between studies, and TL was speculated as one of the possible mechanisms that underlying these fitness effects ((Bouwhuis et al., 2015; Schroeder et al., 2015)Schroeder et al., 2015), however, this is unlikely to be the case in one study (Sparks et al 2022).

Other methodological factors influencing parental age effect on offspring TL estimates included controlling for parental identity and the age of offspring when TL was measured. Not controlling for parental identity resulted in more negative estimates in non-human studies. Since only two studies out of 12 controlled for parental ID, the statistical power of this analysis was limited. However, we still recommend future studies to control for parental identity where possible to reduce the effect of pseudoreplication (Hurlbert, 1984). While in the human analysis we found no effect of offspring age at sampling, in the non-human vertebrate subset, offspring that were measured as juveniles or as juveniles and adults had borderline more negative parental age effects (p=0.05) on offspring telomere length than offspring measured as adults. This might be because adults have undergone more environmental noise exposure which masks the effect of parental age. Future analyses should account for the age at which offspring TL is measured when studying parental age effects. Despite the evidence for the parental environment influence on TL (Lulkiewicz et al., 2020), controlling for parental environment conditions in the study did not affect the offspring TL estimates. This may be because the specific environmental variables included across studies varied. In addition, while some studies included parental environment variables in their models, interactions between parental age and parental environment conditions were not accounted for, which could obscure systematic differences between studies that did and did not control parental environment conditions.

### Time-lag and outcome reporting biases

Significant outcome reporting bias and time-lag bias were detected in human studies, but not in the non-human studies. The asymmetry in the residual-based funnel plot (Figure 8a), together with a significant Egger’s test, indicates that smaller than average effects from studies with small sample sizes and/or low precision underrepresented in human studies. This outcome reporting bias could inflate the overall observed effect. The time-lag bias detected that more recent studies tended to produce effect sizes that were higher than earlier studies. This could be attributed to advancements in research methodologies and technologies (e.g., improvements in the accuracy of TL measurement). Importantly however, this time-lag bias could also be explained by the existing reporting bias favoring large positive parental age effects in human studies. These reporting biases may lead to an overestimation of the parental age effect in human studies, potentially biasing our understanding of its true magnitude. Studies in non-human vertebrates did not detect significant reporting or time-lag bias. This could be because non-human vertebrates have more heterogeneous patterns in terms of parental age effects on offspring TL and thus a reporting bias cannot form, or because there are fewer studies in non-human vertebrates than in humans and therefore we lack of power to detect any biases.

### Implications for future research

Future research should aim to include more non-human vertebrates from a wider variety of taxa, for a better understanding of the parental age effect on offspring TL. For studies in any taxa, we recommend to include multiple longitudinal measurements from the same parents at different ages and separate within- and between-parent effects, and use TRF methods for measuring TL, especially TRF in gel, where possible. Other general suggestions include accounting for cell type, parental identity and offspring age effect in the model. Future studies should account for the effect of offspring sex, especially non-human studies were there is a current knowledge gap. A major limitation of this meta-analysis was the exclusion of many studies due to the lack of effect sizes and standard errors reporting. This highlights the need for standardized reporting practices in primary research to improve data comparability.

## Acknowledgements

We thank members of the Dugdale Research Group for valuable discussion. We thank Simon Verhulst and Agnes Szwarczynska for their invaluable suggestions on the study, as well as Maaike Versteegh for feedback on earlier drafts.

## Data Availability

Datasets and R code used in this study are openly available at:

## Benefits Generated

Benefits from this research accrue from the sharing of our data and results on public databases as described above.

## Author Contributions

HLD, YS and HYJC conceived the study. MV and YS collected the data. MV performed the statistical analyses with input from YS, HYJC and HLD. MV and YS wrote the first draft of the manuscript with input from HYJC and HLD. All authors contributed to the development of the study, provided comments on the manuscript and agreed on the final version of the manuscript to be submitted for publication.

