## Supplemental figures and tables for "Parental age effects on offspring telomere length across vertebrates: a meta-analysis"

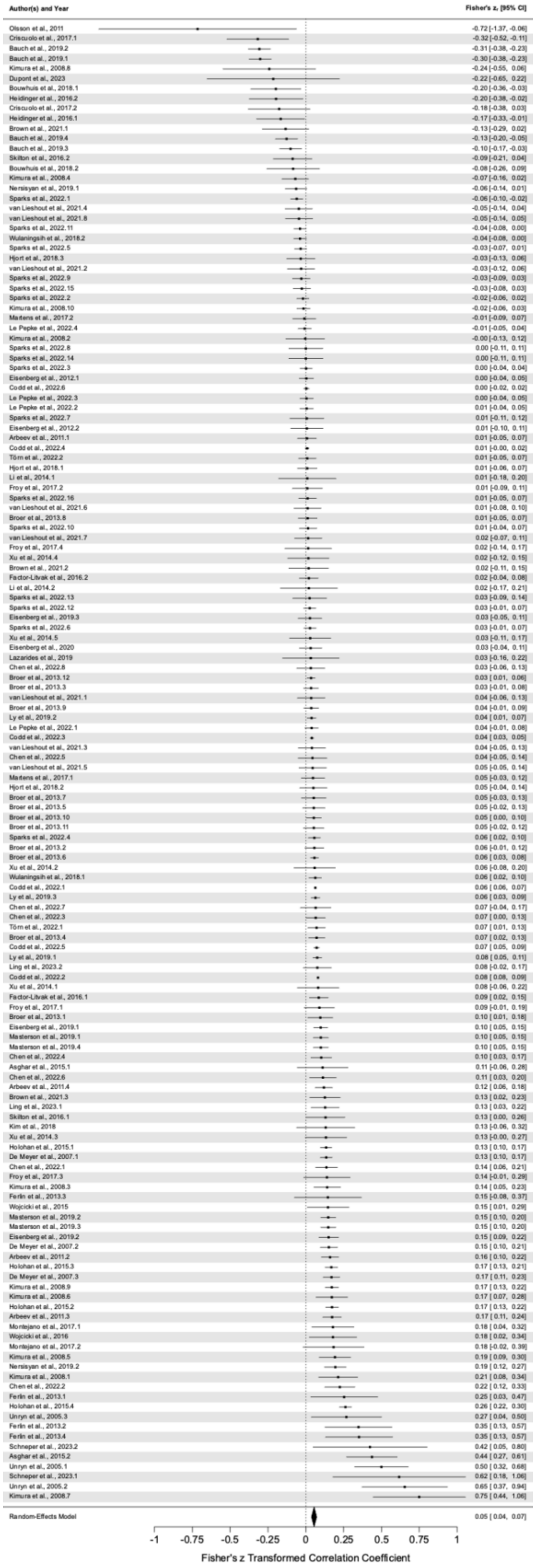
Author(s) and Year
Fisher's z, [95% CI]
Olsson et al., 2011 -0.72 [-1.37, -0.06]
Criscuolo et al., 2017.1 -0.32 [-0.52, -0.11]
Bauch et al., 2019.2 -0.31 [-0.38, -0.23]
Bauch et al., 2019.1 -0.30 [-0.38, -0.23]
Kimura et al., 2008.8 -0.24 [-0.55, 0.06]
Dupont et al., 2023 -0.22 [-0.65, 0.22]
Bouwhuis et al., 2018.1 -0.20 [-0.36, -0.03]
Heidinger et al., 2016.2 -0.20 [-0.38, -0.02]
Criscuolo et al., 2017.2 -0.18 [-0.38, 0.03]
Heidinger et al., 2016.1 -0.17 [-0.33, -0.01]
Brown et al., 2021.1 -0.13 [-0.29, 0.02]
Bauch et al., 2019.4 -0.13 [-0.20, -0.05]
Bauch et al., 2019.3 -0.10 [-0.17, -0.03]
Skilton et al., 2016.2 -0.09 [-0.21, 0.04]
Bouwhuis et al., 2018.2 -0.08 [-0.26, 0.09]
Kimura et al., 2008.4 -0.07 [-0.16, 0.02]
Nersisyan et al., 2019.1 -0.06 [-0.14, 0.01]
Sparks et al., 2022.1 -0.06 [-0.10, -0.02]
van Lieshout et al., 2021.4 -0.05 [-0.14, 0.04]
van Lieshout et al., 2021.8 -0.05 [-0.14, 0.05]
Sparks et al., 2022.11 -0.04 [-0.08, 0.00]
Wulaningsih et al., 2018.2 -0.04 [-0.08, 0.00]
Sparks et al., 2022.5 -0.03 [-0.07, 0.01]
Hjort et al., 2018.3 -0.03 [-0.13, 0.06]
van Lieshout et al., 2021.2 -0.03 [-0.12, 0.06]
Sparks et al., 2022.9 -0.03 [-0.09, 0.03]
Sparks et al., 2022.15 -0.03 [-0.08, 0.03]
Sparks et al., 2022.2 -0.02 [-0.06, 0.02]
Kimura et al., 2008.10 -0.02 [-0.06, 0.03]
Martens et al., 2017.2 -0.01 [-0.09, 0.07]
Le Pepke et al., 2022.4 -0.01 [-0.05, 0.04]
Kimura et al., 2008.2 -0.00 [-0.13, 0.12]
Sparks et al., 2022.8 0.00 [-0.11, 0.11]
Sparks et al., 2022.14 0.00 [-0.11, 0.11]
Sparks et al., 2022.3 0.00 [-0.04, 0.04]
Eisenberg et al., 2012.1 0.00 [-0.04, 0.05]
Codd et al., 2022.6 0.00 [-0.02, 0.02]
Le Pepke et al., 2022.3 0.00 [-0.04, 0.05]
Le Pepke et al., 2022.2 0.01 [-0.04, 0.05]
Sparks et al., 2022.7 0.01 [-0.11, 0.12]
Eisenberg et al., 2012.2 0.01 [-0.10, 0.11]
Arbeev et al., 2011.1 0.01 [-0.05, 0.07]
Codd et al., 2022.4 0.01 [-0.00, 0.02]
Törn et al., 2022.2 0.01 [-0.05, 0.07]
Hjort et al., 2018.1 0.01 [-0.06, 0.07]
Li et al., 2014.1 0.01 [-0.18, 0.20]
Froy et al., 2017.2 0.01 [-0.09, 0.11]
Sparks et al., 2022.16 0.01 [-0.05, 0.07]
van Lieshout et al., 2021.6 0.01 [-0.08, 0.10]
Broer et al., 2013.8 0.01 [-0.05, 0.07]
Sparks et al., 2022.10 0.01 [-0.04, 0.07]
van Lieshout et al., 2021.7 0.02 [-0.07, 0.11]
Froy et al., 2017.4 0.02 [-0.14, 0.17]
Xu et al., 2014.4 0.02 [-0.12, 0.15]
Brown et al., 2021.2 0.02 [-0.11, 0.15]
Factor-Litvak et al., 2016.2 0.02 [-0.04, 0.08]
Li et al., 2014.2 0.02 [-0.17, 0.21]
Sparks et al., 2022.13 0.03 [-0.09, 0.14]
Sparks et al., 2022.12 0.03 [-0.01, 0.07]
Eisenberg et al., 2019.3 0.03 [-0.05, 0.11]
Sparks et al., 2022.6 0.03 [-0.01, 0.07]
Xu et al., 2014.5 0.03 [-0.11, 0.17]
Eisenberg et al., 2020 0.03 [-0.04, 0.11]
Lazarides et al., 2019 0.03 [-0.16, 0.22]
Chen et al., 2022.8 0.03 [-0.06, 0.13]
Broer et al., 2013.12 0.03 [ 0.01, 0.06]
Broer et al., 2013.3 0.03 [-0.01, 0.08]
van Lieshout et al., 2021.1 0.04 [-0.06, 0.13]
Broer et al., 2013.9 0.04 [-0.01, 0.09]
Ly et al., 2019.2 0.04 [ 0.01, 0.07]
Le Pepke et al., 2022.1 0.04 [-0.01, 0.08]
Codd et al., 2022.3 0.04 [ 0.03, 0.05]
van Lieshout et al., 2021.3 0.04 [-0.05, 0.13]
Chen et al., 2022.5 0.04 [-0.05, 0.14]
van Lieshout et al., 2021.5 0.05 [-0.05, 0.14]
Martens et al., 2017.1 0.05 [-0.03, 0.12]
Hjort et al., 2018.2 0.05 [-0.04, 0.14]
Broer et al., 2013.7 0.05 [-0.03, 0.13]
Broer et al., 2013.5 0.05 [-0.02, 0.13]
Broer et al., 2013.10 0.05 [ 0.00, 0.10]
Broer et al., 2013.11 0.05 [-0.02, 0.12]
Sparks et al., 2022.4 0.06 [ 0.02, 0.10]
Broer et al., 2013.2 0.06 [-0.01, 0.12]
Broer et al., 2013.6 0.06 [ 0.03, 0.08]
Xu et al., 2014.2 0.06 [-0.08, 0.20]
Wulaningsih et al., 2018.1 0.06 [ 0.02, 0.10]
Codd et al., 2022.1 0.06 [ 0.06, 0.07]
Ly et al., 2019.3 0.06 [ 0.03, 0.09]
Chen et al., 2022.7 0.07 [-0.04, 0.17]
Chen et al., 2022.3 0.07 [ 0.00, 0.13]
Törn et al., 2022.1 0.07 [ 0.01, 0.13]
Broer et al., 2013.4 0.07 [ 0.02, 0.13]
Codd et al., 2022.5 0.07 [ 0.05, 0.09]
Ly et al., 2019.1 0.08 [ 0.05, 0.11]
Ling et al., 2023.2 0.08 [-0.02, 0.17]
Codd et al., 2022.2 0.08 [ 0.08, 0.09]
Xu et al., 2014.1 0.08 [-0.06, 0.22]
Factor-Litvak et al., 2016.1 0.09 [ 0.02, 0.15]
Froy et al., 2017.1 0.09 [-0.01, 0.19]
Broer et al., 2013.1 0.10 [ 0.01, 0.18]
Eisenberg et al., 2019.1 0.10 [ 0.05, 0.15]
Masterson et al., 2019.1 0.10 [ 0.05, 0.15]
Masterson et al., 2019.4 0.10 [ 0.05, 0.15]
Chen et al., 2022.4 0.10 [ 0.03, 0.17]
Asghar et al., 2015.1 0.11 [-0.06, 0.28]
Chen et al., 2022.6 0.11 [ 0.03, 0.20]
Arbeev et al., 2011.4 0.12 [ 0.06, 0.18]
Brown et al., 2021.3 0.13 [ 0.02, 0.23]
Ling et al., 2023.1 0.13 [ 0.03, 0.22]
Skilton et al., 2016.1 0.13 [ 0.00, 0.26]
Kim et al., 2018 0.13 [-0.06, 0.32]
Xu et al., 2014.3 0.13 [-0.00, 0.27]
Holohan et al., 2015.1 0.13 [ 0.10, 0.17]
De Meyer et al., 2007.1 0.13 [ 0.10, 0.17]
Chen et al., 2022.1 0.14 [ 0.06, 0.21]
Froy et al., 2017.3 0.14 [-0.01, 0.29]
Kimura et al., 2008.3 0.14 [ 0.05, 0.23]
Ferlin et al., 2013.3 0.15 [-0.08, 0.37]
Wojcicki et al., 2015 0.15 [ 0.01, 0.29]
Masterson et al., 2019.2 0.15 [ 0.10, 0.20]
Masterson et al., 2019.3 0.15 [ 0.10, 0.20]
Eisenberg et al., 2019.2 0.15 [ 0.09, 0.22]
De Meyer et al., 2007.2 0.15 [ 0.10, 0.21]
Arbeev et al., 2011.2 0.16 [ 0.10, 0.22]
Holohan et al., 2015.3 0.17 [ 0.13, 0.21]
De Meyer et al., 2007.3 0.17 [ 0.11, 0.23]
Kimura et al., 2008.9 0.17 [ 0.13, 0.22]
Kimura et al., 2008.6 0.17 [ 0.07, 0.28]
Holohan et al., 2015.2 0.17 [ 0.13, 0.22]
Arbeev et al., 2011.3 0.17 [ 0.11, 0.24]
Montejano et al., 2017.1 0.18 [ 0.04, 0.32]
Wojcicki et al., 2016 0.18 [ 0.02, 0.34]
Montejano et al., 2017.2 0.18 [-0.02, 0.39]
Kimura et al., 2008.5 0.19 [ 0.09, 0.30]
Nersisyan et al., 2019.2 0.19 [ 0.12, 0.27]
Kimura et al., 2008.1 0.21 [ 0.08, 0.34]
Chen et al., 2022.2 0.22 [ 0.12, 0.33]
Ferlin et al., 2013.1 0.25 [ 0.03, 0.47]
Holohan et al., 2015.4 0.26 [ 0.22, 0.30]
Unryn et al., 2005.3 0.27 [ 0.04, 0.50]
Ferlin et al., 2013.2 0.35 [ 0.13, 0.57]
Ferlin et al., 2013.4 0.35 [ 0.13, 0.57]
Schneper et al., 2023.2 0.42 [ 0.05, 0.80]
Asghar et al., 2015.2 0.44 [ 0.27, 0.61]
Unryn et al., 2005.1 0.50 [ 0.32, 0.68]
Schneper et al., 2023.1 0.62 [ 0.18, 1.06]
Unryn et al., 2005.2 0.65 [ 0.37, 0.94]
Kimura et al., 2008.7 0.75 [ 0.44, 1.06]
Random-Effects Model 0.05 [ 0.04, 0.07]
-1
-0.75
-0.5
-0.25
0
0.25
0.5
0.75
1
Fisher's z Transformed Correlation Coefficient

**Figure S1**. Non-aggregated forest plot of the individual effect size estimates included in the meta-analysis of parental age at conception effects on offspring telomere length. Each dot represents the estimate effect, with horizontal lines indicating the respective 95% confidence intervals and the dotted vertical line indicating zero.

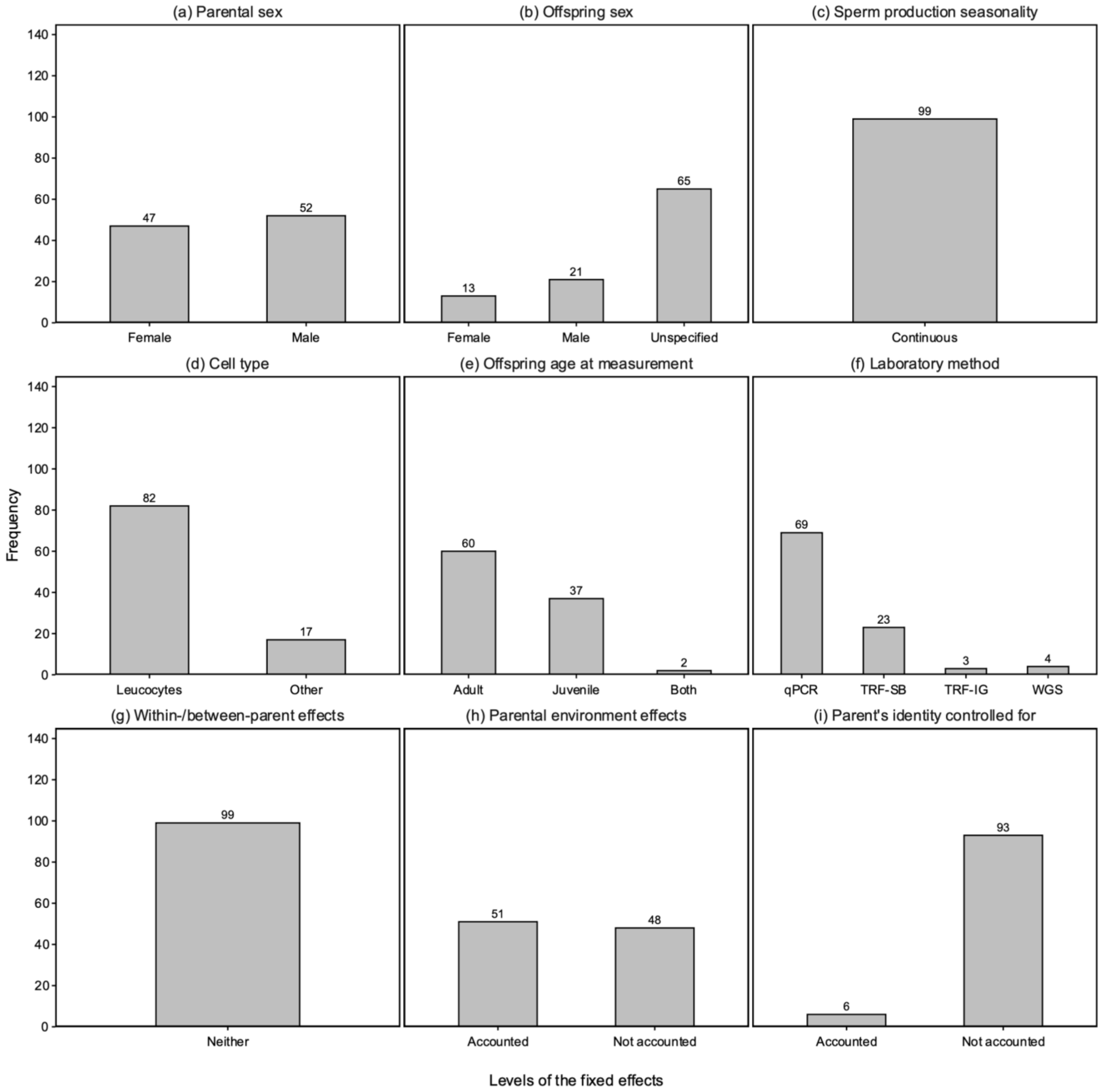

(a) Parental sex
(b) Offspring sex
(c) Sperm production seasonality
(d) Cell type
(e) Offspring age at measurement
(f) Laboratory method
(g) Within-/between-parent effects
(h) Parental environment effects
(i) Parent's identity controlled for
Frequency
0
20
40
60
80
100
120
140
47
52
Female
Male
13
21
65
Female
Male
Unspecified
99
Continuous
82
17
Leucocytes
Other
60
37
2
Adult
Juvenile
Both
69
23
3
4
qPCR
TRF-SB
TRF-IG
WGS
99
Neither
51
48
Accounted
Not accounted
6
93
Accounted
Not accounted
Levels of the fixed effects

**Figure S2.** Bar plots showing the distribution of estimates of parental age effect on offspring telomere lengths across (a) parental sex, (b) offspring sex, (c) sperm production seasonality, (d) cell type from which the telomeres were extracted, (e) offspring age at measurement, (f) laboratory method, (g) whether within-/between-parent effects were accounted for, (h) whether parental environment effects were controlled for, and (i) whether the parent's identity was controlled for in humans. Numbers above the bars represent sample sizes of estimates (N = 99).

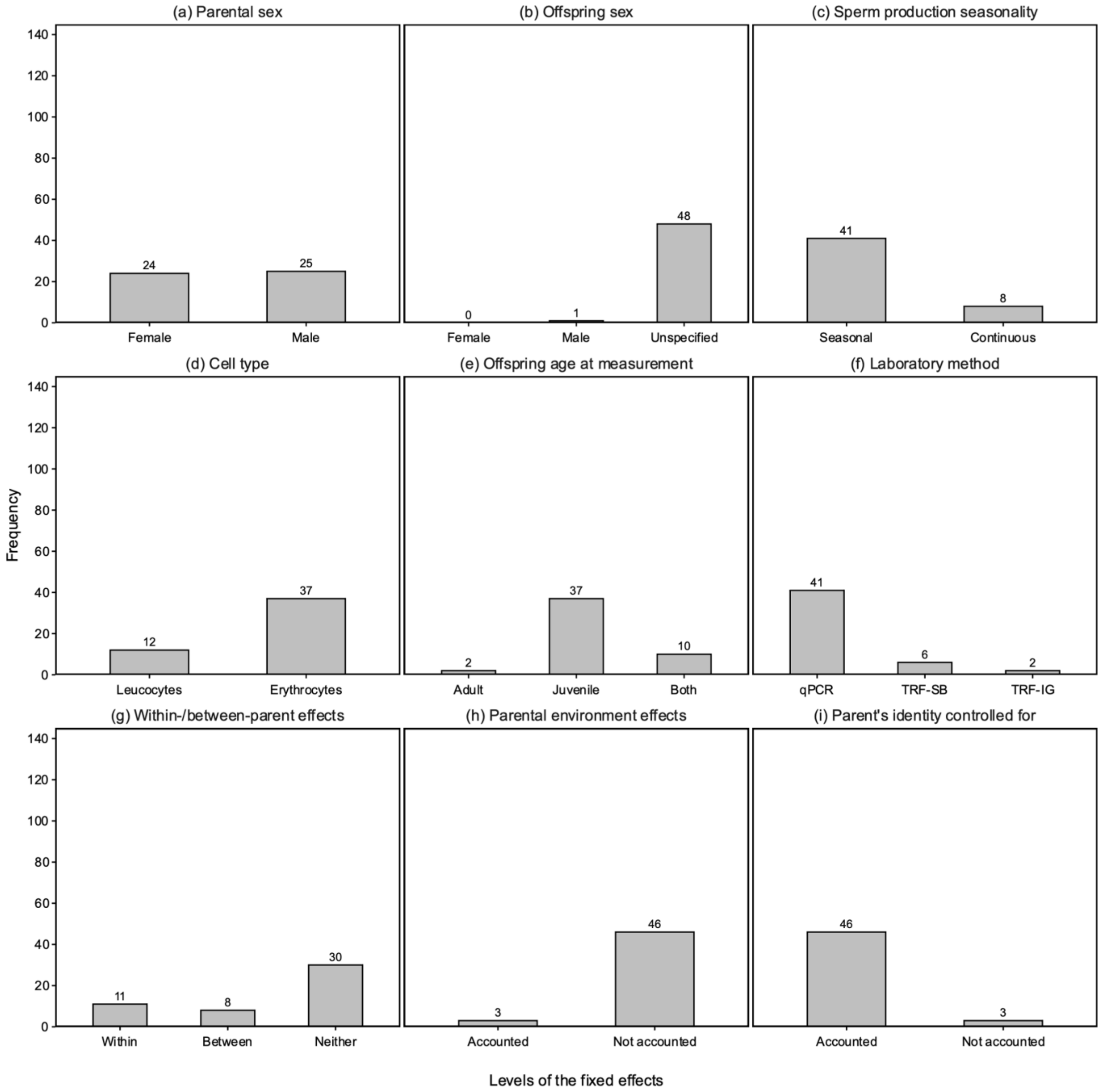

(a) Parental sex
Female
Male
24
25
(b) Offspring sex
Female
Male
Unspecified
0
1
48
(c) Sperm production seasonality
Seasonal
Continuous
41
8
(d) Cell type
Leucocytes
Erythrocytes
12
37
(e) Offspring age at measurement
Adult
Juvenile
Both
2
37
10
(f) Laboratory method
qPCR
TRF-SB
TRF-IG
41
6
2
(g) Within-/between-parent effects
Within
Between
Neither
11
8
30
(h) Parental environment effects
Accounted
Not accounted
3
46
(i) Parent's identity controlled for
Accounted
Not accounted
46
3
Frequency
0
20
40
60
80
100
120
140
Levels of the fixed effects

**Figure S3.** Bar plots showing the distribution of estimates of parental age effect on offspring telomere lengths across (a) parental sex, (b) offspring sex (no study on female offspring was present), (c) sperm production seasonality, (d) cell type from which the telomeres were extracted, (e) offspring age at measurement, (f) laboratory method, (g) whether within-/between-parent effects were accounted for, (h) whether parental environment effects were controlled for, and (i) whether the parent's identity was controlled for in non-human vertebrates. Numbers above the bars represent sample sizes of estimates (N = 49).

**Table S1**. Moderator estimates from the meta-regression models of parental age at conception effect on offspring telomere length in humans (model 3.1) and 12 non-human vertebrates (model 3.2), accounting for study, estimate, species and phylogenetic non-independence and including both interactions (parental and offspring sex, offspring age and telomere tissue). Parentheses in the fixed effects subheadings indicate the reference level; p-values <0.05 are in bold and <0.1 in italics. SE = standard error, 95% CI = confidence interval, DF = degrees of freedom. TRF = terminal restriction fragments, WGS = whole genome sequencing.

| | **Model 3.1 - human subset** | | | **Model 3.2 - non-human vertebrate subset** | | |
|---|---|---|---|---|---|---|
| **Fixed effects** | **estimate (SE)** | **95% CI** | **p-val** | **estimate (SE)** | **95% CI** | **p-val** |
| **intercept** | 0.032 (0.050) | [-0.066, 0.131] | 0.52 | -0.727 (0.360) | [-1.433, -0.022] | **0.04** |
| **Parental sex (Mother)** | | | | | | |
| **Father** | 0.129 (0.031) | [0.068, 0.190] | **< 0.001** | -0.015 (0.014) | [-0.042, 0.012] | 0.28 |
| **Offspring sex (Daughter)** | | | | | | |
| **Son** | 0.114 (0.034) | [0.048, 0.180] | **< 0.001** | | | |
| **Unspecified-sex offspring** | 0.087 (0.030) | [0.028, 0.145] | **0.004** | 0.670 (0.353) | [-0.021, 1.362] | *0.06* |
| **Telomere tissue (other)** | | | | | | |
| **Erythrocytes** | | | | -0.0522 (0.0725) | [-0.1943, 0.0898] | 0.47 |
| **Leukocytes** | -0.064 (0.048) | [-0.157, 0.029] | 0.18 | | | |
| **Sperm production seasonality (Continuous)** | | | | | | |
| **Seasonal** | | | | 0.059 (0.083) | [-0.104, 0.221] | 0.48 |
| **Parental environment effects controlled for (No)** | | | | | | |
| **Present** | 0.003 (0.018) | [-0.032, 0.039] | 0.85 | -0.027 (0.083) | [-0.189, 0.135] | 0.74 |
| **Parent's identity controlled for (No)** | | | | | | |
| **Yes** | -0.023 (0.028) | [-0.078, 0.032] | 0.42 | 0.229 (0.090) | [0.053, 0.405] | **0.011** |
| **Within-/ between-parent effects (Neither)** | | | | | | |
| **Between** | | | | 0.021 (0.024) | [-0.027, 0.069] | 0.40 |
| **Within** | | | | -0.025 (0.024) | [-0.073, 0.023] | 0.30 |
| **Offspring age at measurement (adult)** | | | | | | |
| **Both** | -0.003 (0.039) | [-0.079, 0.074] | 0.94 | -0.167 (0.089) | [-0.342, 0.008] | *0.06* |

| | | | | | | |
|---|---|---|---|---|---|---|
| **Juvenile** | -0.045 (0.053) | [-0.149, 0.060] | 0.40 | -0.158 (0.081) | [-0.315, 0.000] | *0.05* |
| **Laboratory method (qPCR)** | | | | | | |
| **TRF in-gel** | 0.412 (0.073) | [0.268, 0.556] | **< 0.001** | -0.157 (0.088) | [-0.328, 0.015] | *0.07* |
| **TRF Southern blot** | 0.061 (0.019) | [0.023, 0.099] | **0.002** | -0.209 (0.063) | [-0.332, -0.085] | **0.001** |
| **WGS** | 0.0092 (0.0334) | [-0.0563, 0.0747] | 0.78 | | | |
| **Interaction parental sex:offspring sex (mother : daughter)** | | | | | | |
| **Father : Son** | -0.131 (0.043) | [-0.215, -0.047] | **0.002** | | | |
| **Father : unspecified-sex offspring** | -0.104 (0.034) | [-0.170, -0.038] | **0.002** | | | |
| **Interaction telomere tissue : offspring age (other : adult)** | | | | | | |
| **Erythrocytes : Both juvenile and adult** | | | | 0.010 (0.049) | [-0.086, 0.105] | 0.84 |
| **Erythrocytes : Juvenile** | | | | | | |
| **Leucocytes : Juvenile** | 0.018 (0.054) | [-0.087, 0.124] | 0.74 | | | |
| **Random effects** | **Estimate ($\sigma^2$)** | **95% CI** | **Proportion heterogeneity explained** | **Estimate ($\sigma^2$)** | **95% CI** | **Proportion heterogeneity explained** |
| **Study ID** | 0.000 | 0.000, 0.002 | 0.191 | 0.003 | 0.000, 0.018 | 0.767 |
| **Estimate ID** | 0.001 | 0.001, 0.003 | 0.750 | 0.001 | 0.000, 0.003 | 0.200 |
| **Phylogeny** | | | | 0 | 0.000, 0.001 | < 0.001 |
| **$I^2$ total** | 0.941 | | | 0.967 | | |
| **$I^2$ residual** | 0.059 | | | 0.033 | | |
| **Observations (N)** | 99 | | | 49 | | |
| **AIC** | -202.97 | | | -87.20 | | |
| **BIC** | -158.85 | | | -55.04 | | |
| **Log likelihood** | 118.48 | | | 60.60 | | |
| **Deviance** | 226.71 | | | 89.70 | | |
| **DF residual** | 84 | | | 35 | | |

**Table S2.** Post-hoc test estimates and standard errors (SE) from models of parental age at conception effects on offspring telomere length for the laboratory method moderator. TRF = terminal restriction fragments, WGS = whole genome sequencing.

| | **Human studies** | | **Non-human studies** | |
|---|---|---|---|---|
| **Laboratory method** | **estimate** | **SE** | **estimate** | **SE** |
| qPCR | 0.067 | 0.021 | -0.403 | 0.186 |
| TRF in-gel | 0.478 | 0.076 | -0.558 | 0.208 |
| TRF Southern blot | 0.126 | 0.027 | -0.613 | 0.189 |
| WGS | 0.077 | 0.040 | | |

**Table S3.** Post-hoc test results for contrasts between different parental and offspring sex combinations, extracted from the meta-regression model of parental age at conception effects on offspring telomere length in human studies. "F F - M F" represents contrasting the female-parent-female-offspring combination with the male-parent-female-offspring combination; the same applies to the other contrasts, with F = female, M = male, and U = unspecified-sex. SE = standard error. Significant contrasts are in bold.

| contrast | estimate | SE | z ratio | p value |
|---|---|---|---|---|
| **F F - M F** | **-0.128** | **0.031** | **-4.114** | **<0.001** |
| **F F - F U** | **-0.085** | **0.030** | **-2.871** | **0.047** |
| **F F - M U** | **-0.110** | **0.029** | **-3.840** | **0.002** |
| **F F - F M** | **-0.116** | **0.033** | **-3.480** | **0.007** |
| **F F - M M** | **-0.112** | **0.032** | **-3.523** | **0.006** |
| M F - F U | 0.043 | 0.026 | 1.639 | 0.57 |
| M F - M U | 0.018 | 0.024 | 0.751 | 0.98 |
| M F - F M | 0.013 | 0.031 | 0.409 | 0.99 |
| M F - M M | 0.016 | 0.028 | 0.567 | 0.99 |
| F U - M U | -0.024 | 0.013 | -1.825 | 0.45 |
| F U - F M | -0.030 | 0.028 | -1.063 | 0.90 |
| F U - M M | -0.027 | 0.026 | -1.036 | 0.91 |
| M U - F M | -0.006 | 0.027 | -0.208 | 1.00 |
| M U - M M | -0.002 | 0.024 | -0.094 | 1.00 |
| F M - M M | 0.003 | 0.031 | 0.110 | 1.00 |
